# Stem cell quiescence requires PRC2/PRC1-mediated mitochondrial checkpoint

**DOI:** 10.1101/2021.04.21.440825

**Authors:** JR Ishibashi, TH Taslim, AM Hussein, D Brewer, S Liu, S Harper, B Nguyen, J Dang, A Chen, D Del Castillo, J Mathieu, H Ruohola-Baker

## Abstract

Both normal and tumorous stem cells can arrest cell division, avoid apoptosis, and then regenerate lost daughter cells following acute genotoxic insult. This protective, reversible proliferative arrest, known as “quiescence,” is still poorly understood. Here, we show that mTOR-regulated mitophagy is required for radiation insult-induced quiescence in *Drosophila* germline stem cells (GSCs). In GSCs, depletion of mito-fission (Drp1) or mitophagy (Pink1 and Parkin) eliminates entry into quiescence, while depletion of mitochondrial biogenesis (PGC1α) or fusion (Mfn2) eliminates exit from quiescence. We also find that mitophagy-dependent quiescence is under epigenetic control; knockdown of Jarid2 (PRC2) or Pc or Sce (PRC1) stabilizes the mitochondria and locks GSCs out of quiescence, while knockdown of PRC2-specific demethylase, Utx, prevents re-accumulation of the mitochondria and locks GSCs in quiescence. These data suggest that mitochondrial number coordinates reversible quiescence. We further identify that the mechanism of quiescence in both GSCs and human induced pluripotent stem cells (iPSCs) relies on mitophagy to deplete the mitochondrial pool of CycE and limit cell cycle progression. This alternative method of G1/S regulation may present new opportunities for therapeutic purposes.

## INTRODUCTION

Diverse types of stem cells have the capacity to exit cell cycle upon stress, only to reenter under the appropriate conditions; this process, coined quiescence, is distinct from senescence because quiescence can normally be reversed^1,2^. Nutrient-sensitive mechanistic target of rapamycin (mTOR) signaling has been implicated in quiescence, with mTOR activation promoting proliferation and exit from quiescence^3^, and mTOR repression being a hallmark of stem cells in quiescence^1–4^ and diapause^5,6^, with some exceptions^7^. Moreover, quiescence is associated with decreased mitochondrial metabolism and increased macroautophagy^1,2^, hereby referred to as autophagy. Epigenetic remodeling is another hallmark of quiescent stem cell states^1,2,6,8,9^. However, it remains unknown whether there are overarching rules that control entering and exiting quiescence across different types of stem cells.

The germline stem cells (GSCs) of the adult *Drosophila* ovary have been demonstrated to enter into quiescence in response to genotoxic insult, such as gamma irradiation^4,10,11^. GSCs contain an ER-derived organelle called the spectrosome, which anchors the stem cell to the anterior end of the niche and elongates during cell division. A self-renewing GSC division produces a new GSC which remains in the niche, and a cystoblast (CB) which exits the niche and goes on to differentiate further into a germ cell cyst (GCC), and ultimately, into an oocyte. Exposure to insults such as low level irradiation rapidly induces apoptosis in CBs and GCCs, while GSCs survive in a state of quiescence before resuming cell cycle and regenerating lost CBs and GCCs^10^. Prior work in this model of insult-induced quiescence has demonstrated the ability for apoptotic CBs to release a protective ligand, Pvf1, which binds to Tie receptors on GSCs and initiates an antiapoptotic signaling cascade that, through bantam miRNA, represses proapoptotic gene, *Hid*^10^. Additionally, the stress-response transcription factor, FOXO, and the metabolic kinase, mTOR, were shown to be crucial for the entry into and exit from quiescence, respectively^4^. However, the specific cellular processes that FOXO and mTOR regulate in quiescent GSCs are not well characterized^12^.

In this study, we find that mTORC1 core component, *raptor*, is required for GSCs to exit quiescence, and that GATOR1 component, *Nprl3*, and TSC component, *Tsc1*, are each required for GSCs to enter into quiescence. We characterize autophagy as a necessary and sufficient condition for GSC quiescence. We also demonstrate that GSCs require Pink1/Park-mediated mitophagy to enter quiescence and mitochondrial biogenesis and fusion to exit quiescence. We also identify PRC1 and PRC2 as key epigenetic modifiers necessary for both insult-induced mitophagy and quiescence itself, suggesting that epigenetic regulation acts upstream of mitochondrial remodeling. We further show that the mechanism of insult-induced quiescence relies on mitochondrial dynamics to temporally regulate a mitochondrial pool of CycE. We go on to characterize that, not only Drosophila GSCs, but also human iPSCs couple cell cycle progression to mitochondrial count via the mitochondrial reserve of CycE.

## MATERIALS AND METHODS

### Fly stocks and culture conditions

Flies were cultured at 25° C on a cornmeal-yeast-agar-medium supplemented with wet yeast^4,10^. The following stocks were obtained from the Bloomington Drosophila Stock Center at Indiana University: w^1118^ (RRID:<u>BDSC_3605</u>), UAS-Dcr2, w^1118^; nos-Gal4 (RRID:<u>BDSC_25751</u>), w*; UAS-GFP (RRID:<u>BDSC_6874</u>), UAS-Tsc1^RNAi #1^ (RRID:<u>BDSC_35144</u>), UAS-Tsc1^RNAi #2^ (RRID:<u>BDSC_54034</u>), (RRID:<u>BDSC</u>_33914), UAS-Cnc^RNAi^ (RRID:<u>BDSC_</u>40854), UAS-Raptor^RNAi#1^ (RRID:<u>BDSC</u>_41912), UAS-Raptor^RNAi#2^ (RRID:<u>BDSC</u>_34814), UAS-Nprl3^RNAi^ (RRID:<u>BDSC</u>_55384), UAS-Mitf^RNAi#1^ (RRID:<u>BDSC</u>_44561), UAS-Mitf^RNAi#2^ (RRID:<u>BDSC</u>_43998), UAS-Nup44^RNAi^ (RRID:<u>BDSC</u>_39357), UAS-Mio^RNAi^ (RRID:<u>BDSC</u>_57745), UAS-Rictor^RNAi^ (RRID:<u>BDSC</u>_36699), UAS-Rictor^RNAi^ (RRID:<u>BDSC</u>_36584), UAS-Nprl2^RNAi^ RRID:<u>BDSC_57538</u>), UASp-mCherry-Atg8a (RRID:<u>BDSC_37750</u>), UASp-GFP-mCherry-Atg8a (RRID:<u>BDSC_37749</u>), UAS-Atg3^RNAi^ RRID:<u>BDSC_34359</u>), UAS-Atg7^RNAi^ (RRID:<u>BDSC_34369</u>), UAS-Atg5^RNAi^ (RRID:<u>BDSC_34899</u>), UAS-Atg12^RNAi^ RRID:<u>BDSC_34675</u>), UAS-Atg18a^RNAi^ (RRID:<u>BDSC_34714</u>), UAS-Atg14^RNAi^ (RRID:<u>BDSC_40858</u>), UAS-Atg13^RNAi^ (RRID:<u>BDSC_40861</u>), UAS-Atg1^RNAi^ (RRID:<u>BDSC_44034</u>), UAS-Atg16^RNAi^ (RRID:<u>BDSC_58244</u>), UAS-Atg1 (OE-1) (RRID:<u>BDSC</u>_51654), UAS-Atg1B (OE-2) (RRID:<u>BDSC</u>_51655), UAS-ref(2)P^RNAi^ (RRID:<u>BDSC</u>_36111), UAS-RUBCN^RNAi^ (RRID:<u>BDSC</u>_43276), UAS-Atg16^RNAi^ (RRID:<u>BDSC</u>_34358), UAS-Pink1^RNAi^ (RRID:<u>BDSC</u>_38262), UAS-Park^RNAi^ (RRID:<u>BDSC</u>_37509), UAS-Drp1^RNAi^ (RRID:<u>BDSC</u>_51483), UAS-Marf^RNAi^ (RRID:<u>BDSC_55189</u>), UAS-Srl^RNAi^ (RRID:<u>BDSC</u>_33914), UAS-Su(z)12^RNAi^ (RRID:<u>BDSC_33402</u>), UAS-Kdm2^RNAi^ (RRID:<u>BDSC_33699</u>), UAS-trx^RNAi^ (RRID:<u>BDSC_33703</u>), UAS-Pc^RNAi^ (RRID:<u>BDSC_33964</u>), UAS-Gcn5^RNAi^ (RRID:<u>BDSC_33981</u>), UAS-rhi^RNAi^ (RRID:<u>BDSC_34071</u>), UAS-Utx^RNAi^ (RRID:<u>BDSC_34076</u>), UAS-HDAC1^RNAi^ (RRID:<u>BDSC_34846</u>), UAS-Set1^RNAi^ (RRID:<u>BDSC_40931</u>), UAS-mei-41^RNAi^ (RRID:<u>BDSC_41934</u>), UAS-Mt2^RNAi^ (RRID:<u>BDSC_42906</u>), UAS-gpp^RNAi^ (RRID:<u>BDSC_42919</u>), UAS-JIL-1^RNAi^ (RRID:<u>BDSC_57293</u>), UAS-mof^RNAi^ (RRID:<u>BDSC_58281</u>), UAS-Jarid2^RNAi^ (RRID:<u>BDSC_40855</u>) and UAS-Sce^RNAi^ (RRID:<u>BDSC_67924</u>).

### Ionizing radiation treatment

Prior to exposure to gamma-irradiation, 2-4 days old flies (5-6M:15-18F) were kept on cornmeal-yeast-agar-medium augmented with wet yeast for 48 hours at 25°. On the day of irradiation, 2/3 of the females and all males were transferred to empty plastic vials and treated with 50 Gγs of gamma-irradiation. A Cs-137 Mark I Irradiator was used to administer the proper irradiation dosage, according to instructed dosage chart. The remaining 1/3 of the females were not irradiated and were dissected within 1 hour of the others receiving irradiation treatment. After irradiation, the flies were flipped onto a new vial of Standard Diet augmented with wet yeast at 25°. 1/2 of the remaining females were dissected at 1-day post-insult (1dpi). Remaining females were dissected at 2-days post-insult (2dpi) (Fig. S1A).

### H3K27me3 fluorescence intensity quantification

Mean pixel intensity of the H3K27me3 immunofluorescence staining was measured using ImageJ analysis software. Germline Stem Cells were identified by characteristic Adducin staining adjacent to cap cells in each germarium. Regions of Interest (ROI) were drawn around non-dividing GSC nuclei and mean pixel intensity in each ROI was measured inside each ROI. Mean pixel intensity was compared across control, 1-dpi and 2-dpi GSCs.

### Drosophila immunofluorescence analysis

Samples were dissected in cold PBS and then fixed in 4% paraformaldehyde for 15 min at room temperature within 30 min of dissection. Samples were then rinsed in PBT (PBS containing 0.2% Triton X-100), and blocked in PBTB (PBT containing 0.2% BSA, 5% normal goat serum) for at least one hour at room temperature. Samples were stored for up to 72 hours at 4° in PBTB. The following primary antibodies were used: mouse anti-adducin (RRID:<u>AB_528070</u> 1:20), mouse anti-Lamin C (RRID:<u>AB_528339</u> 1:20), mouse anti-ATPsynβ (RRID:<u>AB_301438</u> 1:500), mouse anti-Fasciclin III (RRID:<u>AB_528238</u> 1:50), rat anti-cyclin E (A gift from Helena Richardson, 1:200), rabbit anti-GFP antibody (RRID:AB_221569), mouse anti-mCherry antibody (RRID:<u>AB_11133266</u>), rat anti-Vasa antibody (RRID:AB_760351), rabbit anti-Dcp-1 (RRID:<u>AB_2721060</u> 1:100), and rabbit anti-H3K27me3 (RRID:<u>AB_2561020</u> 1:250). Samples were incubated with primary antibodies for 24 hours at 4°. After washes with PBT, fluorophore-conjugated secondary antibodies were utilized including anti-rabbit Alexa 488 (RRID:<u>AB_221544</u> 1:250), anti-rabbit Alexa 568 (RRID:<u>AB_143157</u> 1:250), anti-mouse Alexa 488 (RRID:<u>AB_2534069</u> 1:250), anti-mouse Alexa 568 (RRID:<u>AB_2535773</u> 1:250), goat antirat 568 antibody (RRID:AB_2534121) and anti-rabbit Alexa 647 (RRID:<u>AB_2535812</u> 1:250), for 1.5–2 hours at room temperature in the dark. Samples then washed with DAPI, diluted with PBT to 2 μm/ml, for 15 minutes to visualize nuclei, followed by two PBT wash. The samples were mounted in mounting medium (21ml of Glycerol, 2.4ml of 10x PBS and 0.468g of N-Propyl Gallate) and analyzed on a Leica SPE5 confocal and Leica SP8 confocal laser-scanning microscope.

### Image Deconvolution and 3D Imaging

Images taken from SPE5 confocal laser-scanning microscope were deconvoluted using Leica LIGHTNING software. 3D hiPSCs images were taken using GE DeltaVision OMX SR super-resolution microscope. Deconvoluted confocal images and OMX images were further analyzed with Imaris (Bitplane) program to construct respective 3D models in Maximum Intensity Projection (MIP) mode for Drosophila GSCs, and Normal Shading for hiPSCs.

### hiPSC culture conditions

The human pluripotent stem cell (hiPSC) line WTC-11 (Coriell Institute, GM25256), previously derived in the Conklin laboratory^13^ and the EGFP-tagged TOMM20 WTC iPSC line generated by the Allen Institute for Cell Science (Coriell Institute, AICS-0011) were cultured on Matrigel growth factor-reduced basement membrane matrix (Corning) in mTeSR media (StemCell Technologies). The cells were treated with Rapamycin (200nM–2μM, Fisher Scientific) or DMSO for indicated periods, and thereafter tested for reversion in the absence of Rapamycin.

### hiPSC gene knockout

Guide RNAs (gRNAs) targeting exon 2 of DRP1, PINK1 and PARKIN (Supplemental Table 2) were designed using <u>CRISPOR.org</u>^14^ and inserted into LentiCRISPRv2 plasmid^15,16^ (a gift from Feng Zhang (Addgene plasmid # 52961), as done before^7^. LentiCRISPRv2 contains two expression cassettes, hSpCas9 and the chimeric guide RNA. The vector was digested using *BsmB*I, and a pair of annealed oligos was cloned into the single guide RNA scaffold. Sequences for the gRNAs and primers used for sequencing can be found in Supplemental Table ST1 and ST2 respectively.

### Viral production

HEK 293FT cells were plated one day before transfection. On the day of transfection, the lentiCRISPRv2 plasmid containing the gRNA was combined with packaging vectors psPAX2 (a gift from Didier Trono, Addgene plasmid # 12260) and pMD2.G (a gift from Didier Trono Addgene plasmid # 12259) in the presence of 1□μg/μL of polyethylenimine (PEI) per 1□μg of DNA. Medium was changed 24□hours later and the lentiviruses were harvested 48 and 72□h after transfection. Viral particles were concentrated using PEG-it (System Biosciences, Inc).

### hiPSCs transduction and selection

hiPSCs (WTC-11) were transduced with lentiCRISPR-v2/gRNA lentiviral particles targeting PINK1, PARKIN or DRP1 in the presence of 4 μg/ml polybrene (Sigma Aldrich). The media was changed the next day. Forty-eight hours after infection, cells were selected with puromycin (1 μg/ml) for two days and genomic DNA was extracted using DNAzol reagent (Invitrogen) according to the manufacturer’s instructions and quantified using Nanodrop ND-1000. Genomic regions flanking the CRISPR target sites were PCR amplified with designed primers (Supplemental Table 3) using GoTaq DNA polymerase (Promega) and sent for Sanger sequencing to determine the insertion and deletion errors generated by CRISPR-Cas9 system in exon 2 of PARKIN, PINK1 and DRP1 genes. The editing efficiency and KO score were determined using the Synthego ICE analysis tool. The mutant lines were analyzed further in this study.

### Cell cycle Analysis via flow cytometry

hiPSC, either WT, or Pink1, Parkin or Drp1 mutants, were seeded in 35 mm dishes at a density of 1.0×10^5^ cells. After 48 h of culturing, cells were treated with 2 μM rapamycin or DMSO and 24 hours later, cells were harvested and cell suspensions were pelleted and washed twice with PBS at 300 g for 5 minutes. Cells were resuspended in 2% FBS in PBS and cells were fixed with 5 mL cold 70% ethanol for at least 24 hours (−20°C) and then centrifuged 10 minutes at 500 x g at 4°C. Cells were washed twice with 3 ml PBS at 400 g for 5 minutes at 4°C and 500 uL PI staining buffer was added (1X PBS, 50 μg/mL PI (Calbiochem), 2μg/mL RNase A (Sigma) and 0.1% Igepal (Sigma)) for 3 hours at 4°C. Cell cycle analysis was performed using the FACS Canto II flowcytometer (BD Biosciences). Data analysis was performed using the FlowJo software (Tree Star, Ashland, OR, USA).

### Protein Extraction and Western Blots Analysis

For protein analysis, 1×10^5^ hiPSCs were plated on 35 mm plates coated with Matrigel. Cells were lysed directly on the plate with a lysis buffer containing 20 mM Tris-HCl pH 7.5, 150 mM NaCl, 15% glycerol, 1% Triton x-100, 1 M β-glycerolphosphate, 0.5 M NaF, 0.1 M sodium pyrophosphate, orthovanadate, PMSF, and 2% sodium dodecyl sulfate (SDS). Twenty-five units of Benzonase^®^ Nuclease (EMD Chemicals, Gibbstown, NJ) was added to the lysis buffer right before use. The protein samples were combined with the 4× Laemmli sample buffer, heated (95°C, 5 min), and run on SDS-PAGE (protean TGX pre-casted 4–20% gradient gel; Bio-rad) and transferred to the nitro-cellulose membrane (Bio-Rad) by semi-dry transfer (Bio-Rad). Membranes were blocked for 1 h with 5% milk or 5% BSA (for antibodies detecting phosphorylated proteins), and incubated in the primary antibodies overnight at 4°C. The antibodies used for western blot were β-actin (Cell Signaling 4970 (1:10000), phospho-mTOR (Ser 2448) (Cell Signaling 5536, 1:1000), mTOR (Cell Signaling 2972, 1:1000), pS6 (Cell Signaling 2215, 1:1000), S6 (Cell Signaling 2117, 1:1000), histone 4 (Lys16) acetylation (H4K16Ac) (Millipore Sigma 07-329, 1:1000), Pink1 (D8G3, Cell Signaling 6946, 1:1000), Parkin (Prk8, Cell Signaling, 4211, 1:1000). The membranes were then incubated with secondary antibodies (1:10000, goat anti-rabbit or goat anti-mouse IgG HRP conjugate (Bio-Rad) for 1 hour and detection was performed using the Immobilon-luminol reagent assay (EMP Millipore).

### Immunofluorescence Staining of hiPSCs

Cells were washed with PBS (2 × 5 min) and then were fixed in 4% paraformaldehyde in PBS for 15 min and blocked for 1 h in 3% BSA+0.1% Triton X-100. The cells were then incubated in primary antibody overnight at 4 °C, washed with PBS (3 × 5 min), incubated with the secondary antibody in 3% BSA+0.1% Triton X-100 for 2 h at room temperature washed (3 × 10 min) and stained with 2 μm/ml DAPI diluted with 1X PBS for 15 min. Mounting media was composed of 21ml of Glycerol, 2.4ml of 10x PBS and 0.468g of N-Propyl Gallate. Analysis was done on a Leica SPE5 Confocal microscope using a ×63 objective and Leica Software. The antibodies used for immunostaining were anti-Oct-4 (Santa Cruz, 1:100), anti-ATP synthase β subunit (Abcam, Ab14730, 1:500), anti-cyclin E human (Santa Cruz, sc-481, 1:100), anti-TFE3 (Sigma Prestige HPA023881, 1:400), anti-HADHA (Abcam, ab54477, 1:250), anti-Phospho-Histone 3 (Ser10) (Millipore Sigma, 06-570, 1:250), Alexa Fluor 568 Phalloidin (Invitrogen, Cat: A12380, 1:300) and Alexa 488- or 647-conjugated secondary antibody (Molecular Probes, 1:250).

### Statistical analysis

All data are presented as the mean of n ≥ 3 experiments with the standard error of the mean (SEM) indicated by error bars, unless otherwise indicated. Statistical significance was determined using chi-squared test (between timepoints) or Student’s t-test (between conditions). Only p values of 0.05 or lower were considered statistically significant (p > 0.05 [ns, not significant], p ≤ 0.05 [*], p ≤ 0.01 [**], p ≤ 0.001 [***], p ≤ 0.0001 [****]). For analysis of quiescence, only fold changes of ≥1.6 were evaluated for significance. Data were compiled and analyzed with Excel for Mac (2020; Microsoft, Seattle, WA, USA).

## RESULTS

### Knockdown of Tsc1 prevents IR-induced quiescence in Drosophila stem cells

After gamma irradiation, FOXO transcription factor is necessary for the entrance into quiescence, and Tor is necessary for the exit from quiescence^4^ (Fig. 1B). Therefore, we sought to characterize the molecular interaction between FOXO and Tor in quiescent GSCs. We first characterized which Tor complex – mTORC1 or mTORC2 – is required for insult-induced quiescence.

**FIGURE 1:**
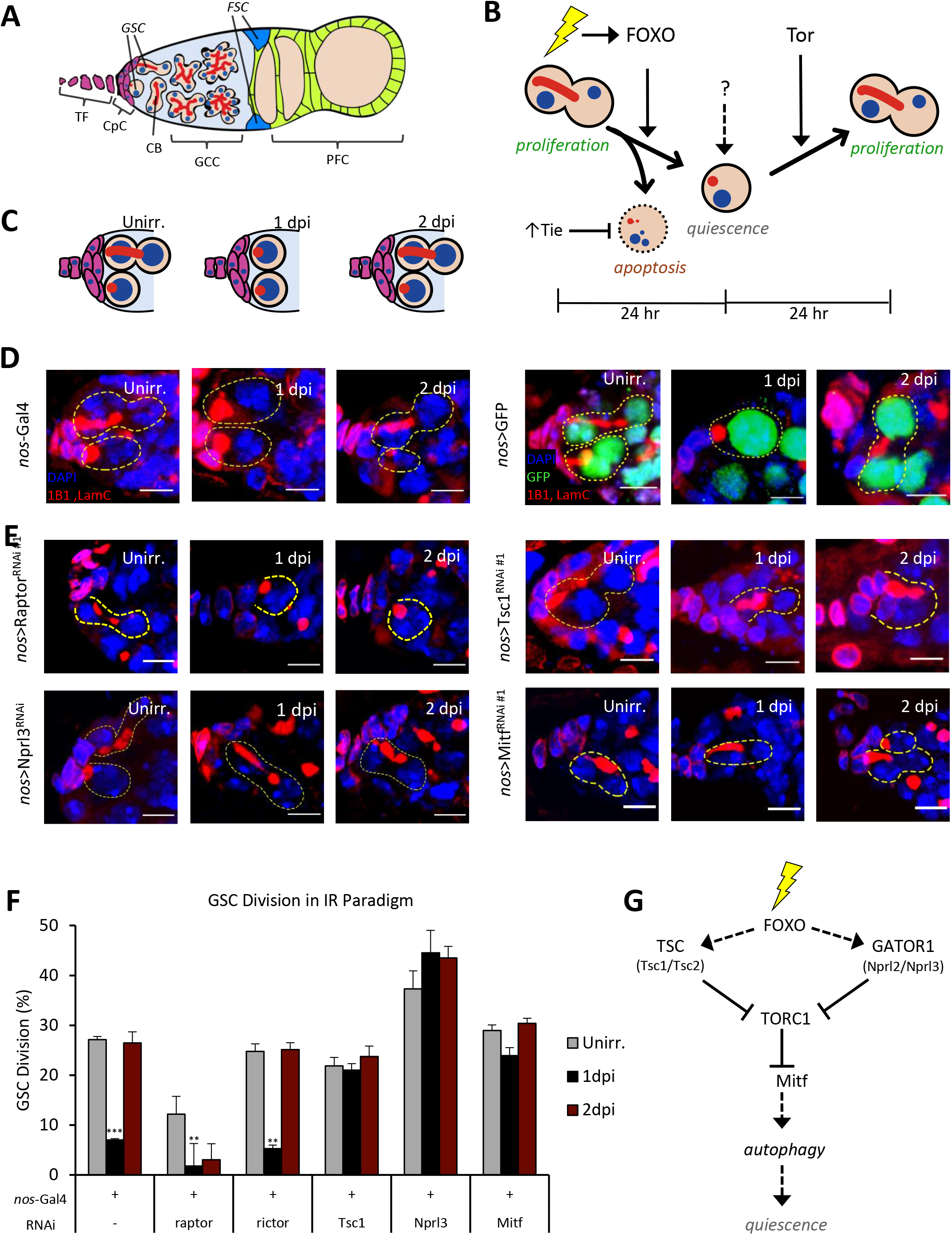
Role of mTORC1 in regulating insult-induced quiescence in female GSCs. **(A)** Representative diagram of a germarium in the Drosophila ovary. TF cells (magenta, cuboidal) comprise the anterior tip of the ovary and connect to the CpCs (magenta, planar); GSCs contain anterior spectrosomes (red, circular) which elongate (red, linear) during division; undifferentiated CBs contain spectrosomes (red, circular) or early fusomes (red, linear) but have no junctions to the CpCs; GCCs are identified by presence of a branched fusome (red, branched); all GSCs and progeny contain nuclei (blue, circular); FSCs (cerulean) can be identified by their distinct triangular morphology; mature GCCs are encapsulated by the PFCs (lime green, planar) and proceed through oogenesis. **(B)** Diagram represents the known role of FOXO and Tor in GSCs, and Tie shown to prevent stem cells from undergoing apoptosis in response to oxidative stress. In the 24 hours following ionizing radiation, FOXO induces quiescence, and in 48 hours following radiation, Tor is needed for the quiescent stem cell to enter the cell cycle. The exact factors which maintain quiescence are poorly understood. **(C)** Cartoon of GSC quiescence. Unirradiated GSCs divide normally, with some containing an elongated spectrosome. 1-day post-insult (1dpi), GSCs arrest cell division and enter quiescence, as evidenced by the absence of elongated spectrosomes. 2 days post-insult (2dpi), GSCs exit quiescence and resume division, as evidenced by the return of elongated spectrosomes. **(D)** Representative confocal microscopy images of nos-Gal4 and nos>GFP from unirradiated, 1dpi, and 2dpi germaria stained with 1B1 (red, spectrosomes/fusomes), LamC (red, Cpc and TF) and DAPI (blue, nuclei), as well as GFP (green, GSCs and progeny). Dotted circle represents GSC (Scale bar 5um). **(E)** Representative confocal microscopy images of listed UAS-RNAi knockdown from unirradiated, 1dpi, and 2dpi germaria stained with 1B1 (red, spectrosomes/fusomes), LamC (red, Cpc and TF) and DAPI (blue, nuclei). Dotted circle represents GSC (Scale bar 5um). **(F)** Percentage of dividing GSCs based on spectrosome elongation. Control GSCs divide at a baseline level when unirradiated (25%), which sharply decreases during quiescence at 1 dpi (4%) and recovers to near baseline rates at 2dpi (14%). In GSC lack of Raptor or Rictor, GSC basal division rate is 12% and 25% respectively, with 24 hours post irradiation drop both division rates to 1% and 5% respectively, and showed that by 48 hours, Raptor KD GSC remains low in division of 3% and Rictor KD GSCs returned to normal division rate of 25%. **(G)** Proposed model by which irradiation activates FOXO, which in turn inhibits mTORC1, possibly by upregulating mTORC1 repressors like the Tuberous Sclerosis Complex (TSC) and GATOR1. mTORC1 inhibition then derepresses Mitf transcription factor, which up regulates target pathway such as autophagy.

In wildtype control flies, before irradiation 27% of GSC show an elongated spectrosome (Fig. 1C,D,F), indicative of active cell division. One day after insult, the GSCs enter into a state of quiescence, as only 5-7% still divide (Fig. 1F). Two days after insult, GSCs resume cell division (26.5%) (Fig. 1F), which means they have successfully exited quiescence. Knockdown (KD) of the mTORC1 component *raptor* by RNAi moderately decreases unirradiated rates of GSC division (12%) (Fig. 1E,F), consistent with mTORC1 activity promoting cell division. These *raptor* KD GSCs (Fig. 1E) fully enter quiescence (2%), but strikingly, GSC division remains low at 2dpi (3%), indicating a failure to exit quiescence (Fig. 1F). In contrast, KD of the mTORC2 component *rictor* by RNAi (Fig. S1B) has no appreciable effect on unirradiated rates of division (25%), and the GSCs properly arrest division at 1dpi (5%) and resume division at 2dpi (25%) (Fig. 1F), similarly to wildtype controls. Taken together, these data show that mTORC1 activation, but not mTORC2, is necessary for the exit from quiescence (Fig. 1G).

To test whether mTORC1 inhibition was necessary for the entry into quiescence, we knocked down components of known mTOR regulatory complexes: TSC, GATOR1, and GATOR2. TSC and GATOR1 are known to inhibit mTOR activity, while GATOR2 inhibits GATOR1 and thereby activates mTORC1^17,18^. When TSC component Tsc1 is knocked down by RNAi in GSCs (Fig. 1E), the rate of GSC division remains mostly unchanged between unirradiated (22%), 1dpi (21%), and 2dpi (24%) (Fig. 1F), suggesting TSC KD eliminates quiescence. Similarly, when GATOR1 component Nprl3 is knocked down (Fig. 1E), the unirradiated rate GSC division is somewhat higher than wildtype (37.5%) (Fig. 1F). However, division remains up at 1dpi (44.5%) and 2dpi (43.5%) (Fig. 1F), suggesting that Nprl3 activity is required to enter quiescence. KD of another GATOR1 component, Nprl2 showed similar inability to arrest cell division (Supplemental Table 1), suggesting that GATOR1-mediated mTORC1 inhibition is essential for insult-induced quiescence (Fig. 1G). Intriguingly, GSCs with GATOR2 components (Nup44a and Mio) knocked down behave very comparably to wildtype controls (Supplemental Table 1), suggesting a nonessential role of GATOR2 in regulating mTORC1 in GSC insult-induced quiescence. These data are consistent with earlier findings that both TSC and GATOR1 become activated in response to the programmed DNA double-stranded break during meiosis in order to mitigate genotoxic stress^18^, and that GATOR2 is not appreciably active in GSCs^19^. From these data, we conclude that mTORC1 repression is necessary to enter into quiescence, and that mTORC1 is repressed by both GATOR1 and TSC activity (Fig. 1G).

Active mTORC1 sequesters lysosomal/autophagy transcription factor Mitf to the cytoplasm by phosphorylation, rendering it inactive^20^. Considering that mTORC1 repression permits Mitf transcription factor activity, we sought to test if it is functional in GSC quiescence. Mitf KD does not dramatically affect unirradiated GSC division rates (29%) (Fig. 1F). However, upon insult, depletion of Mitf abrogates quiescence, as division rates at 1dpi and 2dpi remain high (Fig. 1E-F). Altogether, these data suggest that mTORC1 regulates GSC quiescence through Mitf, a transcription factor that regulates autophagy (Fig. 1G).

### Autophagy-deficient stem cells fail to properly enter into quiescence

Having established that Mitf is required for GSC quiescence after insult, we sought to understand the function of its targets, autophagy genes, in coordinating GSC quiescence. Because autophagy is inhibited by mTORC1, we hypothesized that autophagy may be the key cellular process downstream of mTORC1 signaling.

In canonical autophagy, Atg8a gets incorporated into late autophagosomes and persists in autolysosomes (Fig. 2C). In order to characterize the amount of autophagic degradation before and after radiation insult, we analyzed nos>mCherry-Atg8a GSCs, a reporter of autophagosome/autolysosome formation^21^. We found that before irradiation, a small amount of stem cells (35%) (Fig. 2B, Fig. S2B) contain mCherry-Atg8a punctae, suggesting a low basal amount of autophagy. At 1dpi, many more GSCs contain punctae (87%) which begins to drop back down at 2dpi (54%) (Fig. 2B, Fig. S2B), indicating that autophagosomes and/or autolysosomes are acutely upregulated at 1dpi, concurrent with the GSC quiescence timeline. In order to further characterize the rate of autophagic flux, we analyzed nos>GFP-mCherry-Atg8a GSCs. Unlike mCherry fluorescence, GFP is highly sensitive to pH, and therefore quenches in the low pH of the autolysosome^21^. Using this reporter, autophagosomes are expected to be GFP and mCherry positive, while autolysosomes are expected to be GFP negative but mCherry positive (Fig. S2A). Interestingly, we only observed mCherry positive but GFP negative punctae, suggesting that, in GSCs, mature autophagosomes are rapidly acidified – characteristic of high autophagic flux (Fig. 2A). Furthermore, quantification of the GSCs containing GFP-/mCherry+ punctae gives very similar results to the mCherry-Atg8a construct (Fig. S2C), suggesting little or no contribution of autophagosomes to the mCherry+ punctae. These data align with past results in hematopoietic stem cells suggesting that quiescence is characterized, in part, by the accumulation of autolysosomes^22^.

**FIGURE 2:**
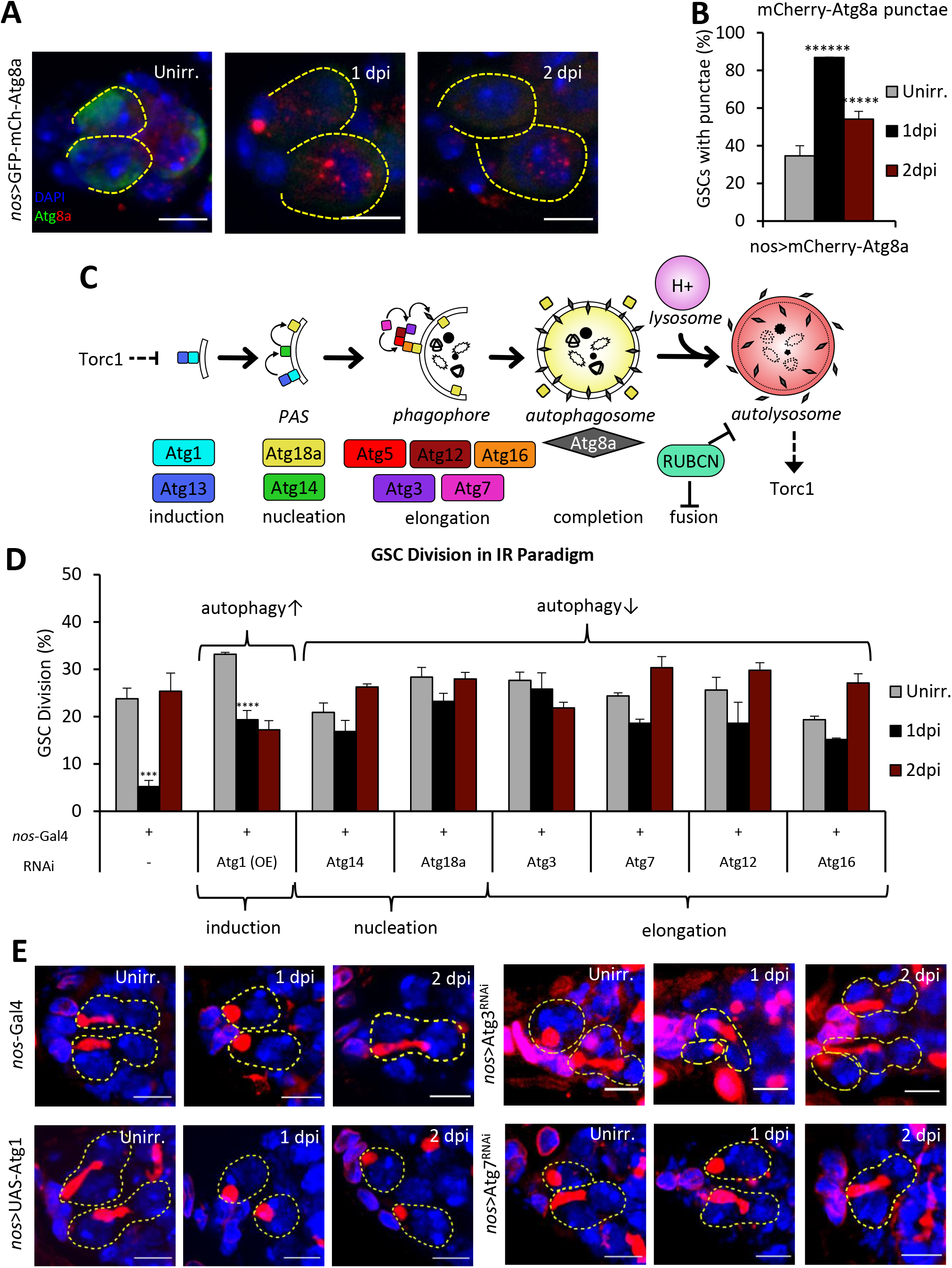
Autophagy-defective germline stem cells display impaired quiescence. **(A)** Representative confocal microscopy images of nos>GFP-mCherry-Atg8a from unirradiated, 1dpi, and 2dpi germaria stained with GFP (green, cytoplasm/autophagosome), mCherry (red, autophagosome/autolysosome), and DAPI (blue, nuclei). Dotted circle represents GSC (Scale bar 5um). **(B)** Quantification of punctae in nos>mCherry-Atg8a germaria, stained with mCherry. A portion of unirradiated GSCs (35%) contain >1 mCherry punctae. At 1dpi, this increases sharply (87%), suggesting acute increase in autophagic degradation. By 2dpi, GSC with punctae decreases (54%), concurrent with the exit from quiescence. **(C)** Schematic of known and hypothesized elements of interplay between autophagy and mTORC1. The Atg1-13 complex initiates autophagy at a double membrane, partly through activation of the Atg14 complex. The Atg14 complex phosphorylates membrane-bound phosphatidylinositol into PI3P. Atg18a localizes to membrane-bound PI3P. Atg7 conjugates Atg12 to Atg5 and Atg16. The Atg12-5-16 complex is recruited to Atg18a on the membrane. Atg7 also conjugates Atg3 to Atg8. Binding of Atg3-8 to Atg12-5-16 activates Atg3, which ligates phosphatidylethanolamine to Atg8. The mature Atg8-PE is lipidated in the membrane, while the other components are excluded from the mature autophagosome. Rubicon is a recently characterized inhibitor of autophagosome maturation. The mature autophagosome then fuses with lysosomes to form the autolysosome, a functional degradative organelle. Successful degradation liberates amino acids, which may serve to activate mTORC1. **(D)** Percentage of dividing GSC with respective autophagy gene RNAi knockdown from unirradiated, 1dpi, and 2dpi germaria. **(E)** Immunofluorescence images of GSC with respective overexpression or knockout of core component of autophagy proteins. Stained with 1B1 (red, spectrosomes/fusomes), LamC (red, Cpc and TF) and DAPI (blue, nuclei). Dotted circle represents GSC (Scale bar 5um).

After confirming that autophagy induction and flux are specifically upregulated after insult, we next sought to characterize whether autophagy was sufficient and/or required for GSC quiescence. Autophagy activation by either Atg1 overexpression (OE) (Fig. 2D-E) or Rubicon KD (Supplemental Table 1) suppressed cell cycle reentry at 2dpi, suggesting that autophagy is at least partly sufficient to prolong quiescence. Similarly, depletion of core autophagy components involved in autophagy induction, phagophore nucleation, membrane elongation, or autophagosome maturation impaired the entrance into quiescence (Fig. 2C-E, Fig. S2D,_Supplemental Table 1). We tested multiple alleles of RNAi against core autophagy components and found comparable results for duplicate alleles (Supplemental Table 1). These data align with findings that mouse embryonic stem cells arrest cell division upon chemical induction of autophagy^23^, suggesting autophagy may be a more universal regulator of stem cell state.

### Mitochondrial remodeling events mediate mTORC1-dependent quiescence

After elucidating the sufficiency and requirement of autophagy for quiescence, we wondered what cellular contents must be degraded by autophagy in order for the germline stem cell to enter into quiescence. Due to the importance of mitochondrial autophagy – or mitophagy – and mitochondrial remodeling in aging and disease^24^, we hypothesized that mitochondrial turnover may be the principle mechanism by which autophagy impairment prohibits GSC quiescence.

We first looked for changes in mitochondrial content and morphology between unirradiated GSCs and recovering GSCs (Fig. 3A-B). Using Vasa to mark the germline and ATP synthase β subunit (ATPsynβ) to image mitochondria, we ascertained that the mitochondrial network of unirradiated GSCs is relatively fused and reticular (90%), with increased mitochondrial density at the interface between the GSC and the cap cells (Fig. 3B), as has been observed before^25^. Strikingly, at 1dpi, the mitochondria were more punctate, with an overall decrease in mitochondrial content (70%; Fig. 3B). Finally, at 2dpi, the mitochondria returned to a more fused network, similar to the mitochondria of unirradiated GSCs (Fig. 3B). These data suggest that mitochondrial turnover coincides with the entry into quiescence, while mitochondrial biogenesis coincides with the exit from quiescence (Fig. 3F). This finding is strikingly similar to findings that proliferating yeast cells contain a tubular meshwork of mitochondria, while quiescent yeast cells have peripheral, fragmented mitochondria^26^, suggesting that mitochondrial morphology controls and/or responds to cell proliferation.

**FIGURE 3:**
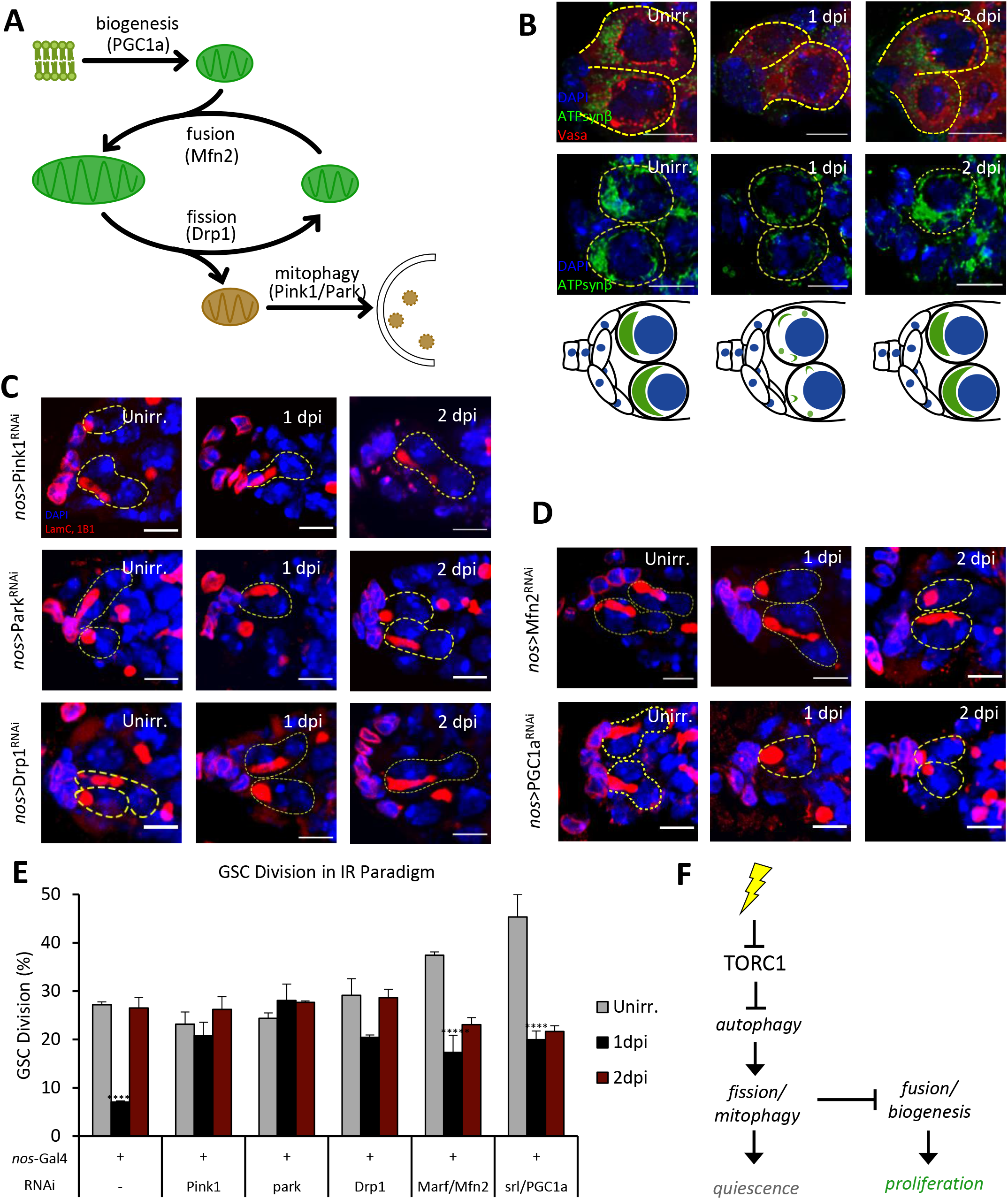
Mitochondrial remodeling events are required for proper coordination of quiescence. **(A)** Current model of mitochondria life cycle based on current literature. **(B)** Representative confocal microscopy images of WT GSCs from unirradiated, 1dpi, and 2dpi germaria stained with VASA (red), ATP synthase β subunit (ATPsynβ) (green, mitochondria), DAPI (blue, nuclei) (upper), or only ATPsynβ and DAPI (lower). (Scale bar 5um). Below is a representative cartoon showing mitochondria fused and polarized towards CpC (magenta) when unirradiated, fragmenting and losing polarization at 1dpi, and refusing and polarizing towards the CpC at 2dpi. **(C)** Representative confocal microscopy images of RNAi knockdown of mitochondrial fission (Drp1) and mitophagy (Pink1/Park) genes from unirradiated, 1dpi, and 2dpi germaria stained with 1B1 (red, spectrosomes/fusomes), LamC (red, Cpc and TF) and DAPI (blue, nuclei). Dotted circle represents GSC (Scale bar 5um). **(D)** Representative confocal microscopy images of RNAi knockdown of mitochondrial fusion (Mfn2) and biogenesis (PGC1α) genes from unirradiated, 1dpi, and 2dpi germaria stained with 1B1 (red, spectrosomes/fusomes), LamC (red, Cpc and TF) and DAPI (blue, nuclei). Dotted circle represents GSC (Scale bar 5um). **(E)** Percentage of dividing GSCs with respective mitochondrial dynamic protein RNAi knockdown from unirradiated, 1dpi, and 2dpi germaria (n = 2 for Drp1, Marf/mfn2, and srl/PGC1α). **(F)** Proposed model where mTORC1 inhibition derepresses autophagy and increases autophagic flux in order to facilitate mitophagy, which is necessary for quiescence. Conversely, mitochondrial fusion/biogenesis is required to exit quiescence and resume proliferation.

We tested if mitophagy is required in GSC quiescence. Pink1 and Park mediate a mitochondrial quality control mechanism that culminates in the clearance of depolarized, dysfunctional mitochondria^27,28^. We show that unirradiated GSCs with Pink1 RNAi KD divide at comparable rates to their wildtype counterparts (Fig. 3E). However, in contrast to controls, Pink1 KD GSCs still divide at 1dpi (21%), failing to enter quiescence (Fig. 3C,E). GSCs with depleted Parkin behave almost identically to Pink KD, where GSCs continue to divide at 1dpi (28%) (Fig. 3C,E). These findings demonstrate that, in addition to autophagy itself, mitophagy is essential for quiescence, and may in fact serve to time quiescence (Fig. 3F).

Due to interaction between mitophagy receptors and mitochondrial dynamics proteins^29,30^, we sought to characterize the roles of canonical regulators of mitochondrial fission, fusion, and biogenesis in GSC quiescence. We first analyzed GTPase Dynamin-related protein, Drp1, which can form helical oligomers that pinch around the outer mitochondrial membrane and induce fission. Drp1 KD GSCs divide normally (29%) before irradiation, but exhibit defects in arresting division at 1dpi (20.5%) (Fig. 3C,E), which implicates mito-fission (Fig. 3A) in the entry into quiescence (Fig. 3F). In contrast, we find that the Marf/mfn2, a GTPase responsible for mitochondrial fusion, and PGC1α, a master transcription factor for mitochondrial biosynthesis, are required for the exit from quiescence. While unirradiated Mfn2-depleted GSCs divide somewhat more than control, they reduce division at 1dpi (>2-fold reduction) but fail to increase cell division by 2dpi (Fig. 3D-E), suggesting a block in the exit from quiescence when fusion is disrupted (Fig. 3F). Almost identically to Mfn2-knockdowns, PGC1α KD GSCs also divide more than wildtype controls and significantly reduce division at 1dpi (>2-fold reduction) but fail to increase division at 2dpi (Fig. 3D-E). These data show that mito-fission and mitophagy regulators promote the entry into quiescence, and mitofusion and biogenesis regulators promote the exit from quiescence (Fig. 3F).

These data show that mitophagy is active and required for insult-induced quiescence, while mitochondrial fusion and biosynthesis are required for the exit from quiescence. These findings suggest that mitochondrial number somehow dictates whether a GSC will divide or be in quiescence.

### Epigenetic modifiers are required for GSC entry and exit from quiescence

Epigenetic proteins modify the chromatin structure to manipulate gene expression, and dysregulated epigenetic modification in cells has been linked to cancer formation^31^. In our tissue-specific RNAi screen we identified multiple epigenetic regulators critical for GSC response to insult (Fig. 4A-B, Fig. S3A), while other epigenetic regulators do not seem to affect the process (Supplemental Table 1). We show that components of the epigenetic repressive machinery of PRC1 and PRC2, Pc/CBX and Sce/RING1 of PRC1, and Jarid2 of PRC2, are required for entry into GSC quiescence, as RNAi knockdown of these proteins abolishes quiescence (Fig. 4A-B, S3A). Interestingly, some transcriptionally-activating epigenetic modifiers were also required for quiescence entry, such as H3K10 kinase, Jil1^32^ and H3K79 methyltransferase, Gpp/DOT1L, and H3K4me2/3 methyltransferase, Set1^33^ (Fig. 4B, Fig. S3A), suggesting a requirement of particular transcriptionally-activating modifications for the entry to quiescence. Additional epigenetic enzymes responsible for recognizing DNA damage, γ-H2Av kinase and mei-41/ATR, regulate entry of quiescence as well (Fig. 4B, Fig. S3A).

**FIGURE 4:**
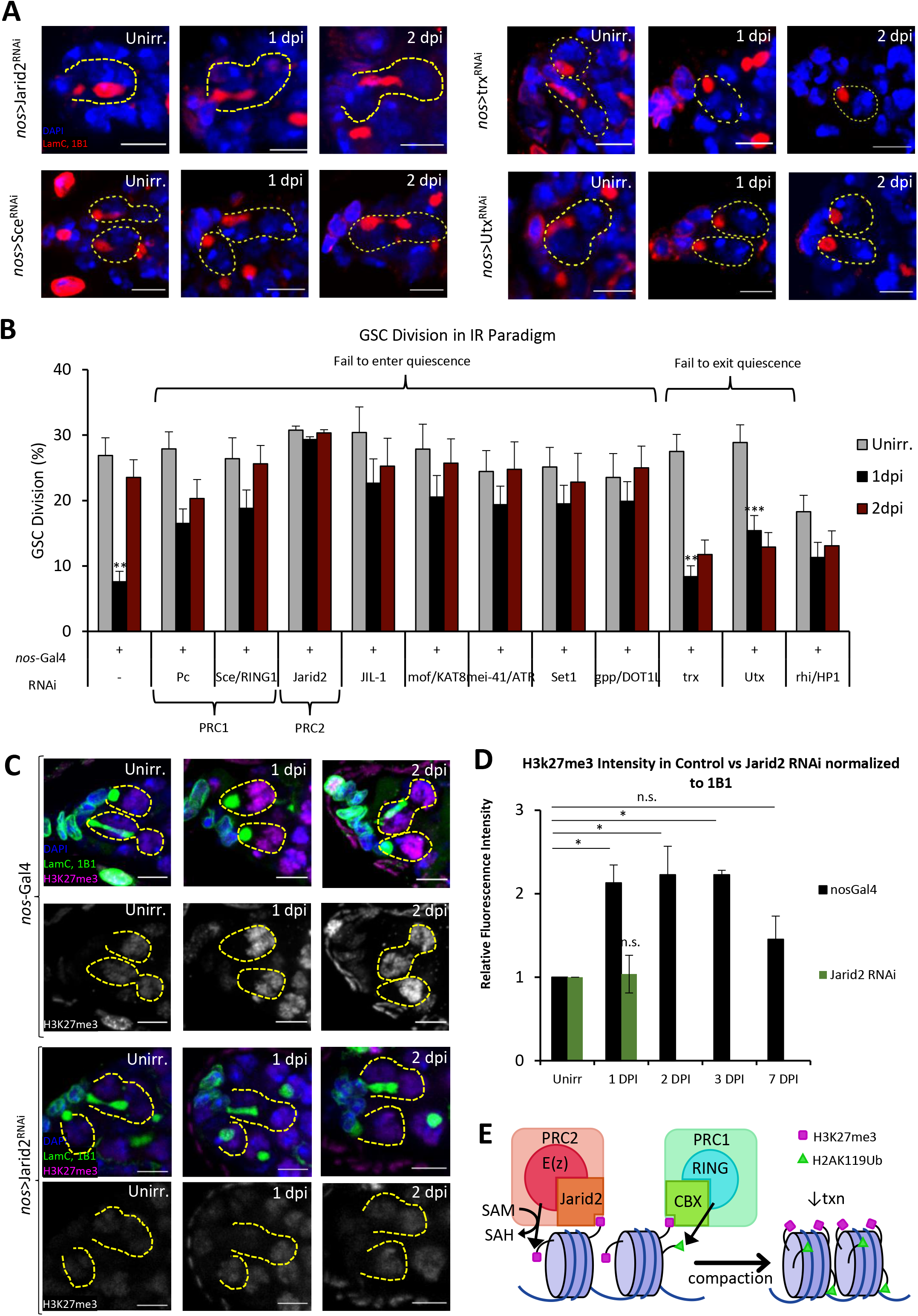
Epigenetic proteins also regulates GSC quiescence. **(A)** Representative confocal microscopy images of RNAi knockdown of various epigenetic regulatory genes from unirradiated, 1dpi, and 2dpi germaria stained with 1B1 (red, spectrosomes/fusomes), LamC (red, Cpc and TF) and DAPI (blue, nuclei). Dotted circle represents GSC (Scale bar 5um). **(B)** Quantification of GSC division. Control GSCs divide at baseline level when unirradiated (27%), which sharply decreases at 1dpi (8%), and recovers to near baseline at 2dpi (24%). Epigenetic proteins like PRC1 components CBX and RING1 are required for entry into quiescence, while other proteins like H3K27me3 demethylase, Utx, are required for the exit from quiescence (n=2 for Sce, JIL-1, mof, mei-41/ATR, rhi/HP1, HDAC1, Kdm2, Gcn5, and Mt2). **(C)** Representative confocal microscopy images of WT and Jarid2-RNAi GSCs from unirradiated, 1dpi, and 2dpi germaria stained with LamC (green, CpC and TF), 1B1 (green, spectrosomes/fusomes), H3K27me3 (magenta) and DAPI (blue, nuclei). Dotted circle represents GSC (Scale bar 5um). **(D)** Quantification of relative fluorescence intensity ofH3K27me3 normalized to 1B1 intensity in WT GSCs in 5 time points and Jarid2 RNAi GSCs in 2 time points. WT H3K27me3 intensity doubles between unirradiated and 1dpi, indicating acute up regulation of H3K27me3 marks upon insult and maintains at 2 dpi and 3 dpi, before decrease seen at 7 dpi. While Jarid2-RNAi GSCs H3k27me3fluorescence intensity remains unchanged throughout the two timepoint. (E) Canonical roles of PRC1 and PRC2 in epigenetic regulation. PRC2 binds to DNA by Jarid2, while E(z) consumes SAM in order to methylate H3K27, leading to transcriptional repression. PRC1 then binds to and recognizes existing H3K27me3 marks through CBX, and then catalyzes monoubiquitination at H2AK119, which further represses transcription.

Conversely, we found another class of epigenetic modifiers that seem to be essential for the exit from quiescence. For instance, RNAi knockdown of Utx, a H3K27me3 demethylase, prevents GSCs from exiting quiescence (Fig. 4A-B). Additional histone modifier, H3K4me1 methyltransferase, Trx^34^, also regulates GSC exit of quiescence, as knockdown of Trx in GSCs causes impairment in the exit of quiescence (Fig. 4A-B). Curiously, rhi, a piRNA pathway component and member of the Heterochromatin protein 1 (HP1) family^35^, likely regulates various aspects of GSC homeostasis and quiescence, as rhi RNAi knockdown causes low baseline division before irradiation and impairs quiescence entry and exit at 1dpi and 2dpi, respectively (Fig. 4B, Fig. S3A).

Because both PRC1 and PRC2 can recognize preexisting H3K27me3 marks (Fig. 4E) and are necessary for entry into quiescence, we tested if H3K27me3 changes throughout GSC quiescence using a H3K27me3 antibody. In wild type GSCs, we detected around 2-fold increase in H3K27me3 mean fluorescence intensity from unirradiated to 1dpi (Fig. 4C,D), which remains high before returning to unirradiated levels by 7dpi (Fig. 4C,D, Fig. S3B). In contrast, when we knockdown PRC2 component Jarid2, no significant increase in H3K27me3 levels is observed after insult (Fig. 4C-D). These results suggest that a global, PRC2 dependent H3K27me3 increase likely directs GSCs to enter quiescence, while removal of site-specific H3K27me3 marks by Utx, but not global H3K27me3 reduction, directs GSCs to exit quiescence.

### Mitophagy-dependent quiescence is under dynamic epigenetic control

Having demonstrated that mitochondrial remodeling and epigenetic regulation are each required for GSC quiescence, we sought to understand whether one might regulate the other in a conserved pathway, as has been shown in other cellular stages^6,8,36,37^. We stained GSCs with ATPsynβ to analyze mitochondrial morphology (Fig. 5A-E, Fig. S4A), and quantified the incidence of fragmented and disorganized mitochondria (Fig. 5F, Fig. S4A). In control flies, only a small portion (5%) of GSCs have fragmented, disorganized mitochondria before irradiation (Fig. 5A,F, Fig. S4A). At 1dpi, the normal anterior mitochondrial network is lost in 70% of the GSCs, and the GSC ratio with dramatically reduced and severely fragmented mitochondria is significantly increased. At 2dpi, the mitochondria show normal morphology and pattern as the GSCs exit quiescence (Fig. 5A,F, Fig. S4A). In contrast, Pink1 KD GSCs, which fail to enter quiescence after insult (Fig. 5B,F) show normal mitochondrial network with very low amounts of fragmentation at 1dpi (3%) (Fig. 5B, Fig. S4A). Interestingly, Sce/RING1 KD GSCs, which fail to enter quiescence after insult (Fig 4A-B), also display low mitochondrial fragmentation (6%) at 1dpi (Fig. 5C,F, Fig. S4A). In contrast, PGC-1 and Utx mutants continue to display segmented and highly reduced mitochondria in 2dpi (Fig. 5D-F). These data show that PRC2 and PRC1 are required for mitochondrial regulated quiescence (Fig. 5G). These data further suggest that mitochondrial remodeling, regulated by epigenetic remodeling, is essential for GSC quiescence.

**FIGURE 5:**
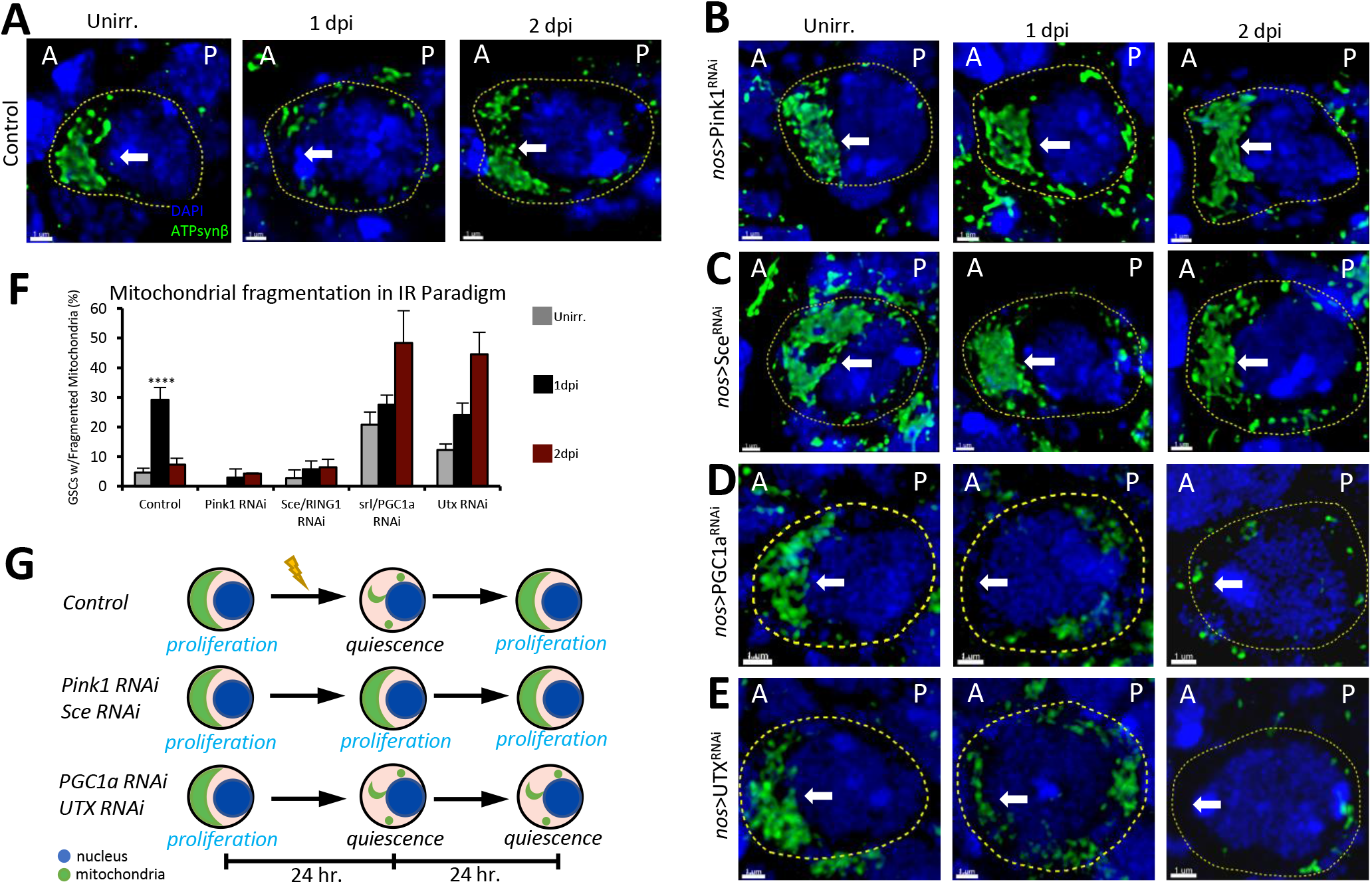
Mitophagy-dependent quiescence is mediated by PRC1. **(A-E)** Representative 3D reconstructed confocal microscopy images of GSCs and their respective knockdowns from unirradiated, 1dpi, and 2dpi stained with DAPI (blue, nuclei) and ATPsynβ (green, mitochondria). (‘A’ denotes anterior side, and ‘P’ denotes the posterior side of the GSC). Arrow points to the area of interest, where mitochondria are typically clustered (Scale bar 1μm). **(F)** Quantification of incidence of fragmented mitochondria where clustered mitochondria are not present in GSC. Images from Fig. S4A were used to generate this quantification. In WT GSCs, there’s very little fragmentation when unirradiated, followed by a sharp accumulation of fragmented mitochondria at 1dpi, which drops back down at 2dpi. Conversely, unirradiated Pink1 RNAi GSCs show no fragmentation when unirradiated, which only marginally increases at 1dpi and 2dpi. Similarly, Sce RNAi GSCs show minimal fragmentation when unirradiated, at 1dpi, and at 2dpi. PGC1a and UTX RNAi GSCs in contrast slightly increased in fragmentation from Unirradiated to 1dpi, and continues to increase in fragmented mitochondria on 2dpi. **(G)** Proposed model by which mitochondria population acts as a checkpoint for the cell cycle state of GSC. Blue represents nucleus and green represents mitochondria.

### Cyclin E localized to the mitochondria is degraded upon genotoxic or chemical insult in GSCs and hiPSCs

The importance of epigenetic regulation of mitophagy further suggests that mitophagy is the metabolic cornerstone of quiescence, as has been recently shown in colorectal cancer^38^. However, the question remains whether mitophagy affects quiescence strictly through bioenergetics, or if mitochondrial number can toggle cell cycle progression through more direct means. Because Drosophila female GSC division doesn’t rely heavily on mitochondrial ATP^39^, we entertained the possibility that the mitochondria played a more direct role in cell cycle progression.

Recent work in mouse fibroblasts and fly ovarian follicle cells has shown that mitochondrial fission/fusion regulate cell cycle through maintenance of a reserve pool of the G1/S regulator Cyclin E (CycE)^40,41^. CycE has also been reported to be targeted for degradation by ubiquitination by Parkin^42^, the E3 ubiquitin ligase required for mitophagy and GSC quiescence (Fig. 3E), and this ubiquitination is Pink1-dependent^43^. Furthermore, Parkin mutations are associated with increased CycE in both cancer and Parkinson’s neurons^44^. Therefore, we hypothesized that Pink1/Parkin-mediated mitophagy might also regulate stem cell quiescence by dictating the amount of available mitochondrial CycE.

We first characterized the CycE subcellular localization in Drosophila GSCs (Fig. 6A). Unirradiated GSCs show striking colocalization of CycE with the mitochondria, as well as the polarized, fused mitochondrial network previously observed. However, one day after insult, the GSCs show a dramatic loss of quantity and polarization of both CycE and mitochondria (Fig. 6A), concurrent with cell cycle arrest or quiescence. Two days after the initial insult, CycE is once again observed colocalizing with the mitochondria, which are once more fused and polarized. Other studies in GSCs have reported an unexplained accumulation of CycE asymmetrically distributed towards the cap cells^45^ that we have now ascertained likely resides on the mitochondria. These data support the hypotheses that mitochondria harbor a pool of CycE in GSCs and that the mitochondrial content dictates cell cycle progression after insult by regulating the CycE availability for G1/S transition.

**FIGURE 6:**
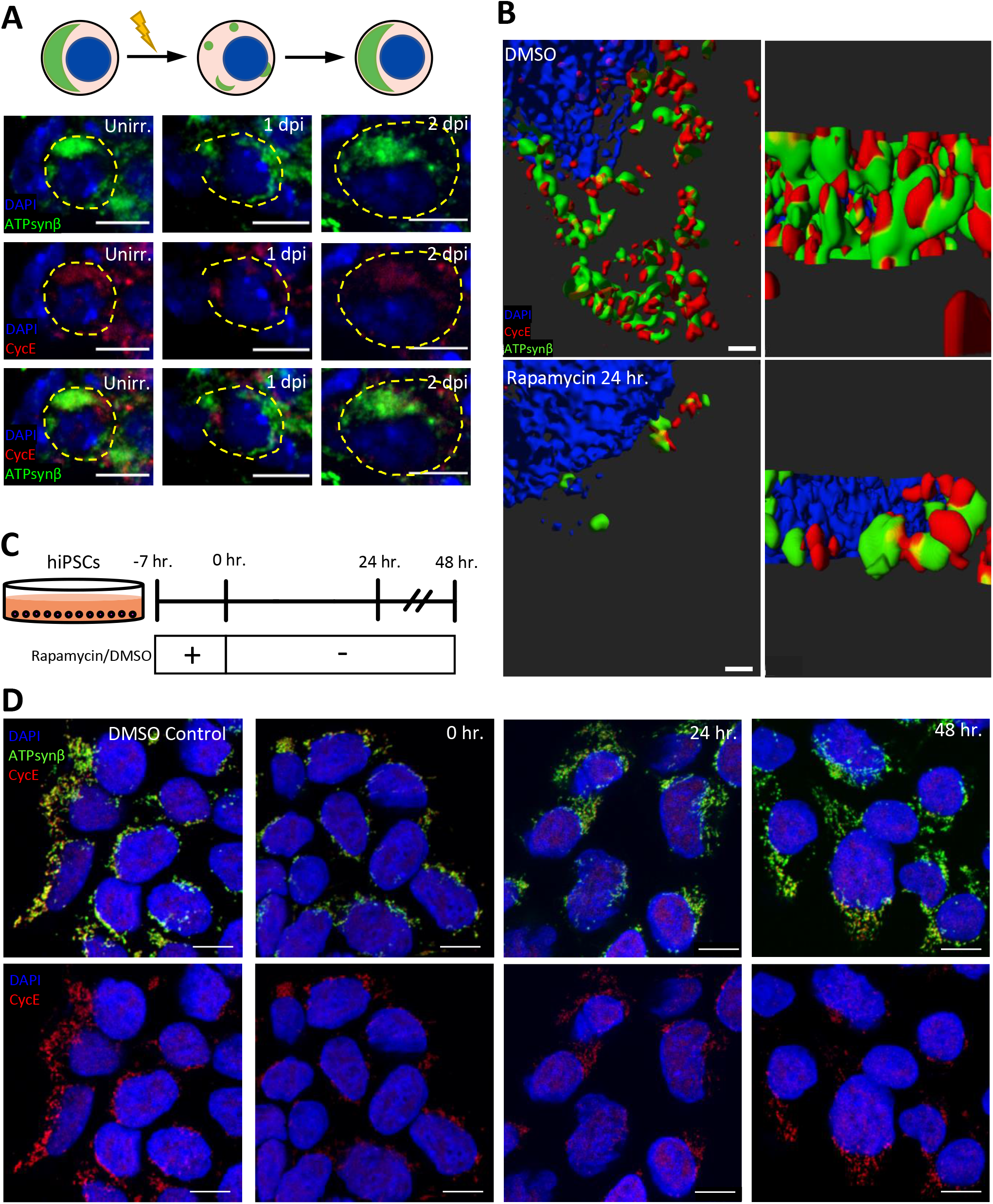
Pool of Cyclin E is observed on Mitochondria in GSCs and iPSCs. **(A)** Representative confocal microscopy images of Drosophila GSCs from unirradiated, 1dpi, and 2dpi germaria stained with ATPsynβ (Mitochondria, green), Cyclin E (red) and DAPI (blue) with cartoon schematic above (Scale bar 5 μm). **(B)** Representative 3D-reconstructed OMX super resolution microscopy images of WTCT cells treated with vehicle control (DMSO) or 2μM rapamycin for 24 hours. Stained with ATPsynβ (mitochondria, green), cyclin E (red) and DAPI (blue) (Scale bar 0.5μm). **(C)** Pulse-chase experiment setup model **(D)** Representative confocal microscopy images of WT iPSCs treated with either vehicle control (DMSO) or rapamycin (2μM) pulse-chase stained with ATPsynβ (mitochondria, green), Cyclin E (red) and DAPI (blue) (Scale bar 10μm). iPSCs shows reduction in both mitochondrial and cyclin E density compared to vehicle control after 7 hours of 2μM rapamycin treatment. = Both mitochondria and Cyclin E density increases after 24 hours, and 48 hours, post rapamycin treatment.

Seeking further confirmation that the number of mitochondria functionally affects CycE, we turned to the surrogate system of human induced pluripotent stem cells (hiPSCs). We used rapamycin, an inhibitor of mTORC1, to induce a diapause-like state in hiPSCS^6^. We show that rapamycin inhibits mTORC1 by western analysis (Fig. S4C). Rapamycin-inhibited hiPSCs show reduced levels of mTORC1 target, pS6, within 7 hours of treatment (Fig. S4C, *left panel*). In addition, we show a stronger inhibition of mTORC1 if rapamycin treatment is extended to 24 hours (Fig. S4C, *right panel*). We also show that this mode of inhibition is reversible as S6 is again phosphorylated within 3 days after mTORC1 reactivation (Fig. S4C). Furthermore, rapamycin treatment results in reversible reduction of epigenetic H4K16Ac mark (Fig. S4C), indicating reduced cellular transcription and a plausible quiescence.

We next analyzed if these quiescent iPSCs show alterations to CycE or mitochondria. We treated iPSCs with 2μM rapamycin to inhibit mTORC1 or vehicle (DMSO) for 24 hours and stained them with CycE and ATPsynβ to analyze CycE and mitochondria subcellular localization (Fig. 6B). By super resolution microscopy, we find that CycE colocalizes with the mitochondria with or without rapamycin treatment (Fig. 6B), confirming that human iPSC mitochondria harbor CycE. Interestingly, upon mTORC1 inhibition by rapamycin, overall CycE levels drop and the mitochondria fragment and degrade, suggesting that rapamycin-induced mitophagy also eliminates mitochondrial CycE (Fig. 6B).

In order to test if mitochondrial fragmentation and CycE depletion can reverse in iPSCs like in GSCs, we treated with rapamycin for 7hr, and then allowed to recover in rapamycin-free medium for up to 48hr (Fig. 6C-D). Control iPSCs show robust mitochondria and CycE (Fig. 6D), but immediately after (0hr) the end of rapamycin treatment, both CycE and mitochondrial number are drastically decreased. After 24-48hr of recovery, the mitochondria and CycE have recovered to how they look the vehicle control. These data confirm that, under conditions of TORC1 inhibition, iPSCs degrade mitochondria and CycE, but that this can be quickly reversed.

### Mitophagy dictates quiescence in human iPSCs

To further investigate the importance of mitophagy in regulating the stability of CycE, we generated hiPSC line with CRISPR induced mutations in the mitochondrial serine/threonine protein kinase, Pink1. Pink1 contains N-terminal mitochondrial localization sequence, transmembrane sequence, Ser/Thr kinase domain and C-terminal regulatory domain^46^. We engineered the CRISPR/Cas9/guide system to induce mutations at or prior to the kinase domain, thereby eliminating the protein’s normal functions (Fig. 7A). Accordingly, using lentivirus-mediated CRISPR-Cas9, 90% of the cells generated had indels near the PAM region (Fig. 7B). We generated CRISPR mutants in two genetic backgrounds and showed that in both cases the mutations had dramatically reduced the level of Pink1 protein (Fig. 7B).

**FIGURE 7:**
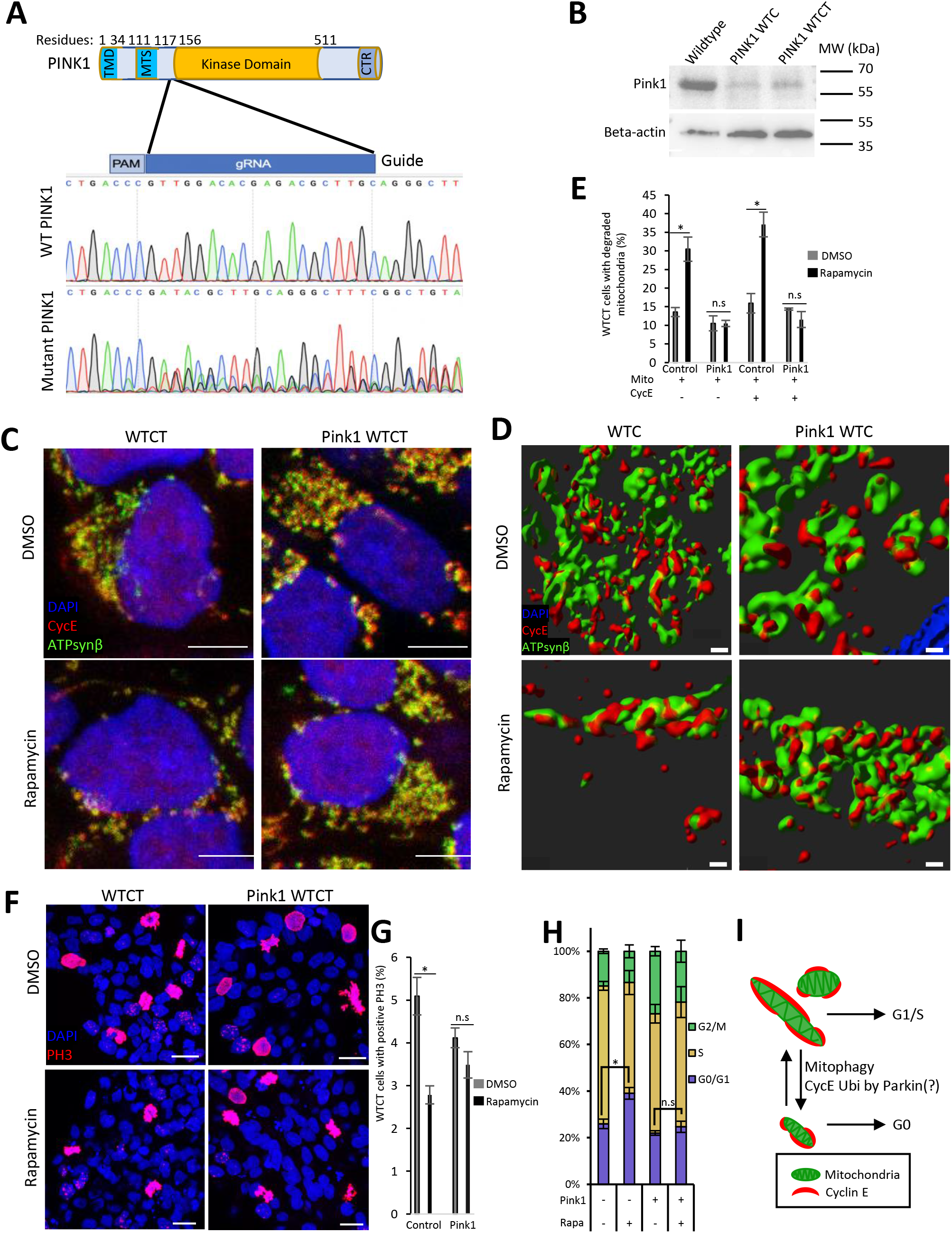
Mitophagy in iPSCs controls both mitochondrial Cyclin E and cell cycle. **(A)** Pink1 structure with guide RNA location indicated and DNA sequencing chromatogram comparing wildtype (WT) Pink1 to mutant Pink1, showing a mixed pool of mutants and a loss of the wildtype sequence. **(B)** Western Blot showing lysates from Control WTC-11, Pink1 mutant WTC and Pink1 mutant WTCT, showing Pink1 protein knocked down in both mutant pools when compared to control **(C)** Representative confocal microscopy images of wildtype WTCT or Pink1 mutant WTCT treated with either vehicle control (DMSO) or rapamycin (2μM) stained with DAPI (blue), CycE (red) and ATPsynβ (mitochondria, green) (Scale bar 5μm). **(D)** Representative 3D reconstructed OMX super resolution microscopy images of wildtype WTC or Pink1 mutant WTC stained with DAPI (blue), CycE (red), and ATPsynβ (mitochondria, green) (Scale bar 0.2μm). **(E)** Quantification of mitochondrial degradation in WTCT vs Pink1 mutant treated with either vehicle control (DMSO) or rapamycin (2μM), suggesting that Pink1 mutants can’t degrade their mitochondria in response rapamycin. **(F)** Representative confocal microscopy images of wildtype WTCT or Pink1 mutant WTCT treated with either vehicle control (DMSO) or rapamycin (2μM) stained with DAPI (blue), PH3 (proliferating cell nuclei, red) (Scale bar 15μm). **(G)** Quantification of PH3 incidence in WTCT vs Pink1 mutant treated with either vehicle control (DMSO) or rapamycin (2μM), suggesting that Pink1 mutants fail to halt cell cycle progression as efficiently as wildtype. **(H)** Quantification of FACS analysis of cell cycle by propidium iodide staining in WTC or Pink1KD treated with DMSO or rapamycin (2μM). **(I)** Model of proposed mechanism in which mitochondria can normally stabilize CycE and promote G1-S transition, while mitophagy induction will reduce CycE – perhaps through direct ubiquitination by Parkin – and keep the stem cells in quiescence/G0.

Using immunofluorescence microscopy, we treated both control iPSC and Pink1 mutant iPSC with Rapamycin or vehicle for 24 hours and observed a dramatic change in the cellular behavior. As before (Fig. 6B,D), control iPSCs show dense, fused mitochondrial networks, with only (~10%) of cells containing reduced mitochondria and CycE, but after 24 hour treatment with 2μM rapamycin mitochondrial and CycE levels reduced significantly (~27-35%) (Fig. 7C-E; Fig. S5A). In contrast, Pink1 mutant iPSCs showed low (~5%) mitochondrial and CycE reduction regardless of whether they were treated with DMSO or rapamycin, suggesting that Pink1/Parkin-mediated mitophagy is required for CycE degradation (Fig. 7C-E; Fig. S5A). Using PH3 to assess stem cell proliferation, we also found that, in control iPSCs, roughly 50% fewer cells were dividing after rapamycin treatment (Fig. 7F-G, Fig. S5B,F). Compared to control, Pink1 mutant iPSCs show less cell cycle arrest, with only 29% fewer stem cells still dividing (Fig. 7F-G Fig. S5B,F).

In addition to microscopic analysis, we also stained iPSC and Pink1 mutant iPSC with PI for cell cycle analysis by flow cytometry. Control iPSCs treated with vehicle (DMSO) for 24hr show a portion of cells in G0/G1 (25.5%), a large portion in S (58.4%), and the remainder in G2/M (14.7%) (Fig. 7H, Fig. S5G). Conversely, after 24hr of rapamycin treatment, iPSCs show significant G0/G1 arrest (38.7%), consistent with the idea that rapamycin induces a diapause-like state of cellular quiescence in human iPSCs. Remarkably, rapamycin treated Pink1 mutant iPSCs show no significant difference when compared to vehicle treated Pink1 mutant iPSCs. Moreover, neither vehicle-nor rapamycin-treated Pink1 mutant iPSCs differ significantly from control iPSCs (Fig. 7H, Fig. S5G). The inability for Pink1 mutant iPSCs to arrest in G0/G1 is consistent with mitophagy reducing CycE and therefore G1-S transition. These data indicate that hiPSCs, like Drosophila GSCs, utilize a noncanonical method of regulating the available reservoir of CycE via Pink1/Parkin-mediated mitophagy. Together, these findings across stem cell types suggest that diverse stem cells may rely on mitochondrial count to regulate available CycE, and consequently stem cell quiescence.

## DISCUSSION

In this study, we show that quiescence in Drosophila germline stem cells relies on the amount of mitochondria; failure to fragment and degrade mitochondria abolishes quiescence, allowing cell cycle to continue, while failure to synthesize and fuse mitochondria impairs the exit from quiescence. We further show that mitochondrial number is regulated by mTORC1 and PRC1/2, and most importantly, mechanistically regulates cell cycle by serving as a harbor for CycE (Fig. 7I).

We show that GSC quiescence is mTORC1-dependent and largely mTORC2-independent. We further characterized that GSCs require both TSC and GATOR complexes, canonical mTORC1 repressors, to arrest proliferation, as depletion of components of either complex abrogates quiescence. Having characterized some key molecules upstream of mTORC1, we then showed that mTOR-responsive transcription factor, Mitf^20^, is required for quiescence. In humans, TFE3 has been shown normally to be phosphorylated by mTOR, sequestered out of the nucleus and rendered inactive^7,9^. However, when mTOR is inactivated, TFE3 stays in the nucleus and induces transcription of its context-dependent downstream targets, including autophagy regulators. The requirement of Mitf, orthologous to human MITF, TFE3, and TFEB, implicates upregulation of autophagosome formation and lysosome acidification in GSC quiescence^20,47^. We show that autophagy is acutely induced following irradiation, as evidenced by an accumulation of mCherry-Atg8a+ punctae, and furthermore that autophagy is required for quiescence, as depletion of autophagy genes abolishes stem cell quiescence. In addition, we showed that autophagy GOF through either Atg1 OE or RUBCN KD is sufficient to suppress the exit from quiescence. Autophagy has been shown to be necessary for stem cell quiescence in some contexts^48–50^, and necessary for proliferation in other contexts^51,52^. Our data unequivocally shows that, in Drosophila female germline stem cells, autophagy is necessary and sufficient for quiescence.

In addition to the general requirement of autophagy for quiescence, we sought to characterize key targets of autophagic degradation. We found that the normally fused mitochondrial network of germline stem cells fragments after irradiation, and that network re-fusion occurs concurrently with the return to proliferation. We further showed that mitophagy effectors, Pink and Parkin, as well as mito-fission protein Drp1, are each necessary for quiescence, consistent with mito-fission being a prerequisite for mitochondrial degradation^25^. These data support the hypothesis that autophagy regulates quiescence primarily by facilitating mitophagy. In the context of regeneration perinatal mammalian cardiomyocytes have been shown to undergo a critical Parkin-dependent mitophagy event that similarly coincides with mTOR inhibition and loss of regenerative capacity^53^. Conversely, we show mitochondrial biogenesis transcription factor, PGC1α, and mito-fusion protein, Mfn2, are required to properly exit from quiescence. In some cases mTORC1 repression has been shown to increase mito-fusion by reducing Drp1 activity^54^, so it is plausible that temporary mTORC1 repression during quiescence stimulates mito-fusion in a negative feedback loop to reactivate mTORC1 and eventually end quiescence. Future research is needed to understand whether manipulating signaling between mTOR and mitochondria could grant regenerative capacity to nonregenerative tissues like the adult heart. Furthermore, it will be important to dissect in detail if and how mitochondrial dynamics play a role in GSC age-related senescence, as mitochondria have shown to play a critical role in senescence in many cell types^56,57,58,59,60,61^.

After characterizing the mitochondrial requirements for entering and exiting quiescence in female GSCs, we also demonstrate that specific epigenetic proteins regulate stem cell quiescence. Previous studies have shown that Polycomb group proteins (PcG) are required in many adult stem cells to maintain stemness. In satellite stem cells, expression of H3K27me3 help promote stemness and self-renewal^62–64^, Downregulation of PcG proteins in satellite cells led to impairment of self-renewal and loss of quiescent satellite stem cell pool^65^. In hematopoietic stem and progenitor cells (HSPCs), PRC1 component Ring1 allows self-renewal^66^. Here, we show that PcG proteins Pc/CBX and Sce/Ring1 in PRC1 are required for GSCs to enter insult-induced reversible quiescence, which alleviates replicative stress and allow DNA damage repair. Similar to our results, Pc homolog Cbx7 in African killifish was recently shown to be required for maintaining diapause, which is a temporal pause of proliferative development^9^. Moreover, we show that H3K27me3 is upregulated in a JARID2-dependent manner upon insult and that Utx/Kdm6a, H3K27me3 demethylase, is required to exit quiescence. Altogether, these data show involvement of PcG proteins in regulation of stem cell quiescence. he first time that PRC1/PRC2 complex activity is essential for the regulation of insult-induced quiescence in GSCs. Interestingly, we also find that PRC1 and PRC2 are each required for mitochondrial fragmentation and degradation, suggesting that epigenetic remodeling is required for mitophagy in the first place. Further work is needed to identify the key target genes regulated in this manner.

Considering the requirement of mitochondrial degradation for entry into quiescence and the reciprocal requirement of mitochondrial fusion and biogenesis for exit from quiescence, we argue that the number of mitochondria serve as the final gatekeeper when entering and exiting quiescence. Previous studies have revealed that mitochondrial metabolites can rate-limit epigenetic remodeling events^6,8,36,37^. Furthermore, in the hematopoietic system, autophagy has been shown to affect mitochondria and thereby epigenetic metabolite-regulated epigenetic marks^49^. Since we have demonstrated that GSCs require mitophagy to enter quiescence, it is unlikely that mitochondrial metabolism is highly active during quiescence. Therefore, it is plausible that once PRC1/PRC2 upregulate mitophagy in order to enter quiescence, low mitochondrial count helps maintain the quiescent epigenome in part by retrograde signaling back to the nucleus. The tricarboxylic acid cycle (TCA) cycle supplies citrate and alpha-ketoglutarate, substrates for activating epigenetic modifications - histone acetylation and histone/DNA demethylation^67^. It is therefore possible that, in addition to controlling CycE, mitophagy also reduces TCA metabolites, thereby disabling activating histone modifications that would otherwise impair cell cycle arrest. This hypothesis is further supported by our finding that mitochondrial biogenesis is required to exit quiescence, suggesting that in regeneration, increased TCA cycle could sufficiently increase alpha-ketoglutarate enough to allow demethylases like Utx/Kdm6a to use it as a cofactor to erase repressive H3K27me3 marks at specific genes and reenter cell cycle. It’s plausible mitophagy could further affect quiescence by limiting the extent of contact between the mitochondria and nucleus and dampening retrograde signaling^68^. It will be important to investigate if similar epigenetic regulation of mitochondrial dynamics controls insult-induced quiescence in other stem cell types and whether senescence in aging stem cells can be reversed by altering mitochondrial dynamics.

The two stem cell types discussed in this paper, insulted Drosophila GSCs and human iPSCs, share an atypical cell cycle regulation in that both are refractory to the typical G1/S phase checkpoint regulators, CDKIs^4,69–72^. Lack of CDKI inhibitor function in iPSC was recently demonstrated by findings revealing that absence of CDKIs increases the iPSC reprogramming efficiency^69^. Moreover, both cell types can enter quiescence during normal development. Insulted GSC enter quiescence within one day of insult and exit quiescence within two days, while PSC can enter a quiescent stage called diapause for an undetermined, but controlled period. In both cases, once conditions are again favorable, the quiescent cell can re-enter cell cycle and regenerate the tissue. This is a particularly extraordinary detour for pluripotent stem cells in the blastocyst, as the zygote would normally proceed from fertilization, through implantation and embryogenesis, to fetal development without halt. However, as neither stem cell type relies on the canonical CDKIs to inhibit CycE, the regulator of stem cell quiescence remains at-large. We have now identified an alternative mode: mitochondrial count, which in turn stabilizes the critical G1/S cell cycle regulator, CycE. We show that mitochondrial count regulates whether the stem cell divides or not. When mitochondrial number is reduced by mitophagy, CycE levels drop and the stem cell halts before G1/S transition, entering a reversible quiescence. Notably Pink1 knockdown eliminates cell cycle arrest in response to rapamycin, revealing that mitochondrial remodeling actively regulates CycE in iPSC quiescence. It will be important to interrogate which stem cell types can utilize this alternative method of G1/S control, and whether this phenomenon can be leveraged for therapeutic purposes.

## Acknowledgments

We thank Prof. Helena Richardson for kindly providing Drosophila CycE Abs. We would like to thank Filippo Artoni, Arianne Caudal, Ondina Palmeira, Ellen Ward, Xiaosheng Yang, Blair Zhao, and other members of the Ruohola-Baker lab for their stimulating discussion and valuable comments. We thank Lynn & Mike Garvey Imaging Core at the University of Washington and the Cell Analysis Facility Flow Cytometry Shared Resource Lab in the Department of Immunology at the University of Washington. This work is supported by the Biological Mechanisms of Healthy Aging Training Program NIH T32AG066574 for A.M.H and support from Hahn Family and partly by grants from the National Institutes of Health R01GM097372, R01GM083867, 1P01GM081619, U01HL099997; UO1HL099993 for H.R.-B.

## SUPPLEMENTAL FIGURE LEGENDS

**SUPPLEMENTAL FIGURE 1: Role of mTORC1 in regulating insult-induced quiescence in female GSCs**

**(A)** Experimental set up for irradiation model; 1–2-week-old flies are placed on standard food supplemented with fresh wet yeast paste two days before irradiation. On the day of irradiation, 1/3 of females are dissected without irradiation, while the other 2/3 and males are treated with 50 Gys ionizing radiation. One day post insult, 1/2 of the remaining females are dissecting. At two days post insult, the remaining females are dissected and males are sacrificed. **(B)** Representative confocal microscopy images of Rictor or Nrf2 RNAi knockdown from unirradiated, 1dpi, and 2dpi germaria stained with 1B1 (red, spectrosomes/fusomes), LamC (red, Cpc and TF) and DAPI (blue, nuclei). Dotted circle represents GSC (Scale bar 5μm).

**SUPPLEMENTAL FIGURE 2: Autophagy-defective germline stem cells display impaired quiescence**

**(A)** Schematic diagram of how UASp-EGFP-mCherry-Atg8a tandem fusion can be used to study autophagic flux. **(B)** Representative confocal microscropy images of nos>mCherry-Atg8a from unirradiated, 1dpi, and 2dpi germaria stained with mCherry (red, autophagosome/autolysosome), VASA (cyan) and DAPI (blue, nuclei). Dotted circle represents GSC (Scale bar 5μm). **(C)** Bar graph depicting the proportion of punctae+ GSCs/total GSCs in nos>GFP-mCherry-Atg8a. **(D)** Immunofluorescence images of GSCs with respective core autophagy component RNAi knockdown. Stained with 1B1 (red, spectrosomes/fusomes), LamC (red, Cpc and TF) and DAPI (blue, nuclei). Dotted circle represents GSC (Scale bar 5μm).

**SUPPLEMENTAL FIGURE 3: Epigenetic proteins regulate GSC quiescence**

**(A)** Representative confocal microscopy images of epigenetic genes RNAi knockdown from unirradiated, 1dpi, and 2dpi germaria stained with 1B1 (red, spectrosomes/fusomes), LamC (red, Cpc and TF) and DAPI (blue, nuclei). Dotted circle represents GSC (Scale bar 5μm). **(B)** Representative confocal microscopy images of WT GSCs from 3dpi and 7dpi germaria stained with LamC (green, CpC and TF), 1B1 (green, spectrosomes/fusomes), H3K27me3 (magenta) and DAPI (blue, nuclei). Dotted circle represents GSC (Scale bar 5μm).

**SUPPLEMENTAL FIGURE 4: Mitochondrial degradation is required for GSCs to enter quiescence**

**(A)** Representative confocal microscopy images of GSCs with respective knockdowns from unirradiated, 1dpi, and 2dpi germarias, which were used for fragmentation quantification on Fig. 5F. stained with DAPI (blue, nuclei) and ATPsynβ (green, mitochondria) throughout the insult timepoint (Scale bar 5μm). (B) WTCT cells treated with 2μM rapamycin for 3 hours, 7 hours or 24 hours, stained with ATPsynβ (mitochondria, red), Cyclin E (green) and DAPI (blue) (Scale bar 5μm). **(C)** Shows the western blot for pulse chase experiment of WT iPSCs treated with either vehicle control (DMSO) or rapamycin (2μM) for 7 hours (left panel) or 24 hours (right panel) followed by reversion (normal iPSCs growth media) for another 96 hours. pmTOR, pS6 and H4k16Ac level is seen to be low by 7 or 24 hours post 2μM rapamycin treatment and their level increases back drastically more than control by 96 hours post reversion. (D) Representative high magnification 3D reconstructed OMX super resolution microscopy images of WTCT cells treated with DMSO vehicle or 2μM rapamycin for 24 hours, stained with ATPsynβ (mitochondria, green), Cyclin E (red) and DAPI (blue) (Scale bar 0.1 μm) **(E)** Summary model of Drosophila GSC showing an initial pules of FOXO inhibits mTORC1, which in turn derepresses fission, mitophagy, and PRC1. Ultimately, mitochondrial fragmentation and degradation seem to constitute a “checkpoint” of sorts before entering quiescence. Then, in order to exit quiescence, H3K27me3 demethylase needs to reactivate specific genes, likely those involved in mitochondrial fusion and biogenesis.

**SUPPLEMENTAL FIGURE 5: iPSCs require mitophagy to regulate Cyclin E**

**(A-D)** Representative confocal microscopy images of control and Pink1-KD iPSCs, treated with either DMSO or 2μM rapamycin for 24 hours. **(A)** Control and Pink1-KD WTCT cells stained with ATPsynβ (mitochondria, green), Cyclin E (red) and DAPI (blue) (Scale bar 15μm). **(B)** Control and Pink1-KD WTC cells stained with PH3 (mitosis marker, red), and DAPI (blue) (Scale bar 15μm). (C-D) Deconvoluted images taken from SP8 confocal microscope. **(C)** Control and Pink1-KD WTC cells stained with ATPsynβ (mitochondria, green), Cyclin E (red) and DAPI (blue) (Scale bar 10μm) **(D)** High magnification of control and Pink1-KD WTC cells stained with ATPsynβ (mitochondria, green), Cyclin E (red) and DAPI (blue) (Scale bar 5μm). **(E)** Quantification of cells without a dense cluster of mitochondria in control vs Pink1-KD WTC treated with control DMSO or 2μM rapamycin for 24 hours. One group of control and Pink1 (left half) is quantified based on the mitochondria, the other group (right half) includes the criteria of only quantifying mitochondrial-CycE positive cells. Each group shows control WTC increase in cells without dense cluster of mitochondria when treated with rapamycin vs DMSO, whereas Pink1-KD WTC cells maintains their dense cluster of mitochondria when treated with rapamycin. Grey bar shows quantification for DMSO treated cells and black bar for 2μM rapamycin treated cells. Data is from duplicate experiments. **(F)** Quantification of PH3 positive cells in control vs Pink1-KD WTC treated with DMSO control vs 2μM rapamycin for 24 hours. Control WTC shows decrease in positive PH3 cells by half when treated with rapamycin compared to DMSO control, whereas Pink1-KD WTC shows slightly higher PH3 positive cells when treated with rapamycin compared to control WTC cells. n=3. (G) Representative FACS cell cycle analysis traces from WTC or Pink1 mutant WTC treated with vehicle (DMSO) or rapamycin (2μM) for 24 hours.

**SUPPLEMENTAL TABLE 1: Excel file compiling all IR paradigm source data relevant to the experiments.**

Data are organized into sheets that they represent in the main figure, followed by genotype, time points and replicate number.

**SUPPLEMENTAL TABLE 2:**
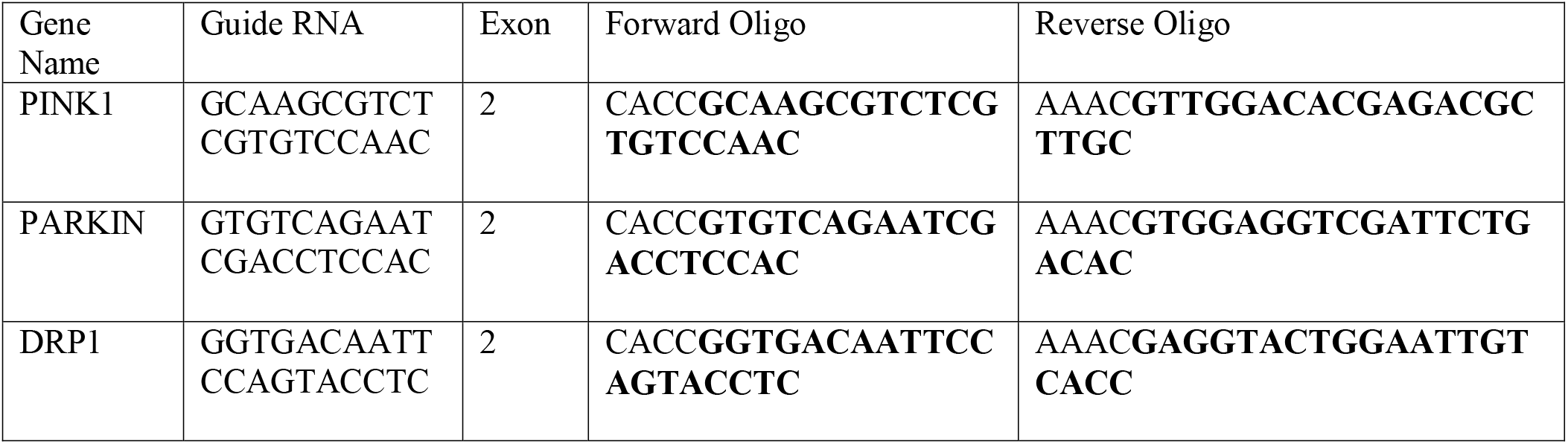
Guide RNAs for PINK1, PARKIN and DRP1 genes

**SUPPLEMENTAL TABLE 3:**
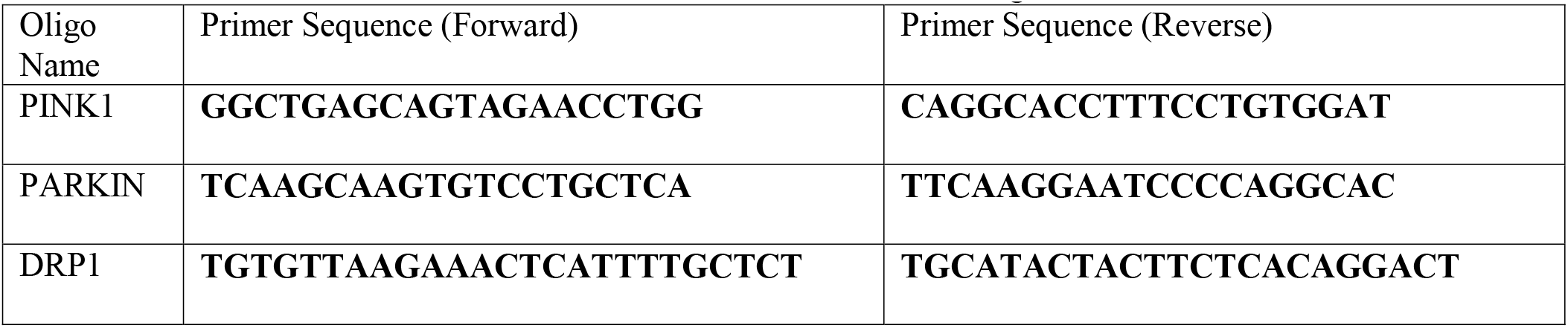
Primers for PINK1, PARKIN and DRP1 genes

## Notes

### Competing Interest Statement

The authors have declared no competing interest.

