## Supplementary figures and images for "Stem cell quiescence requires PRC2/PRC1-mediated mitochondrial checkpoint"

### Supplemental Figures

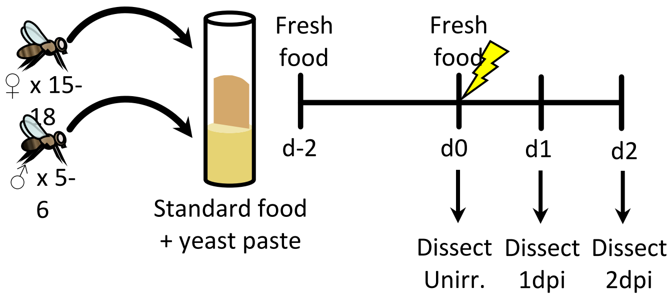

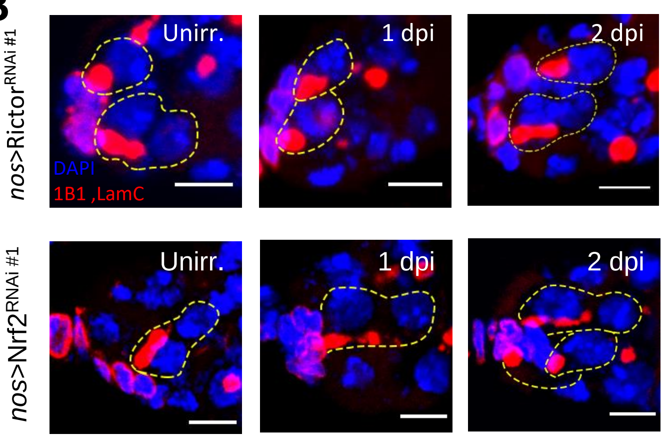

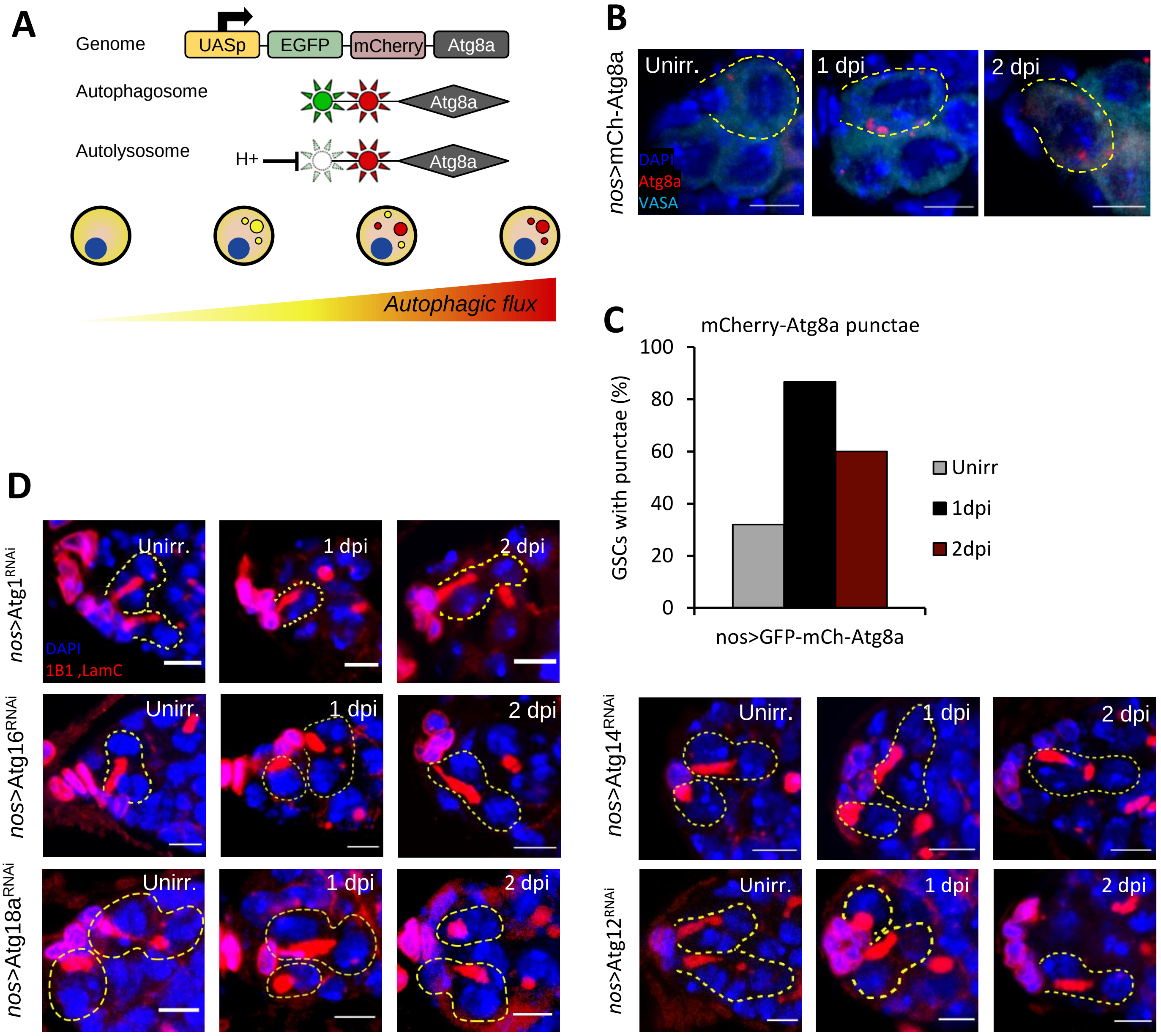

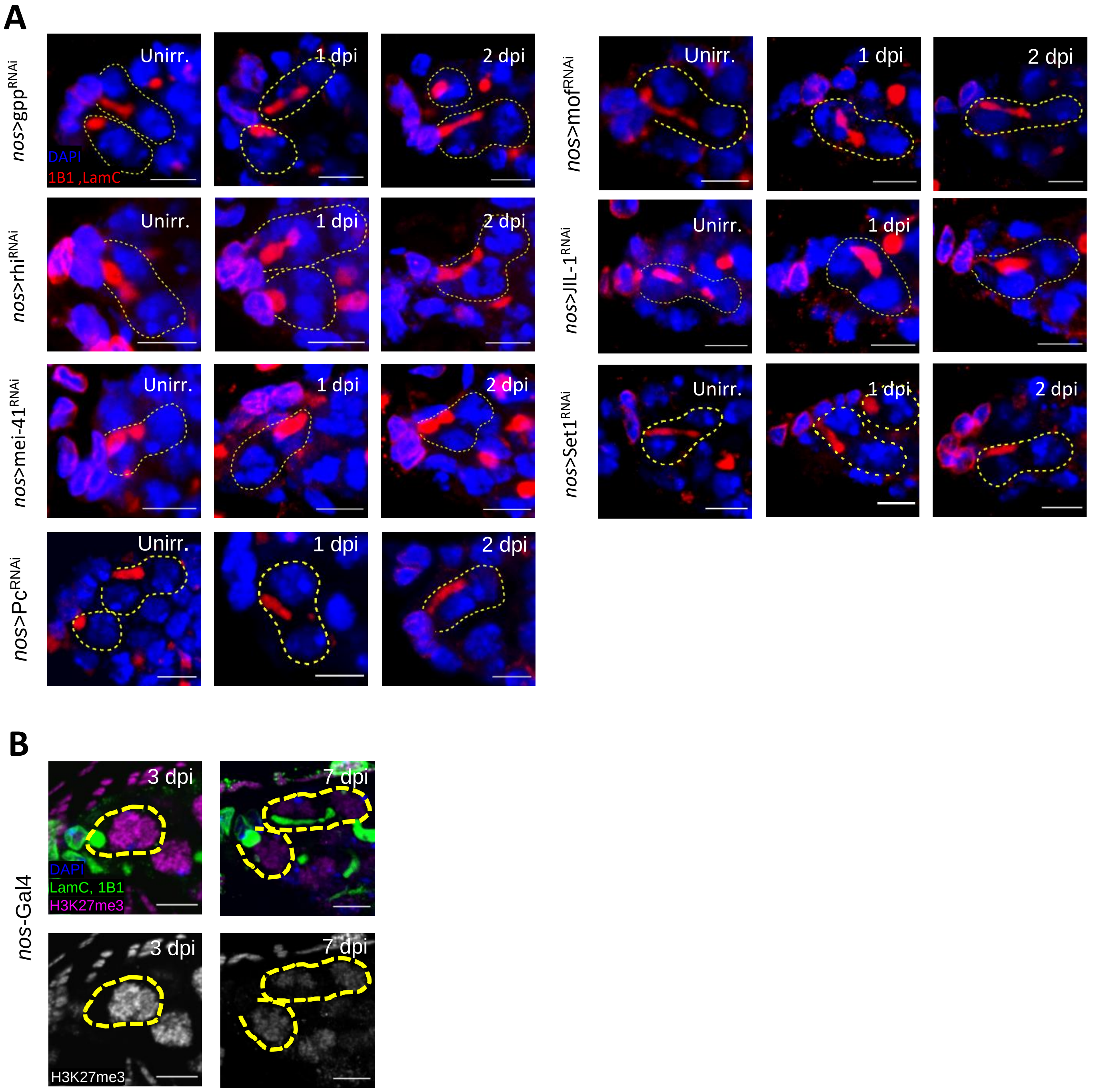

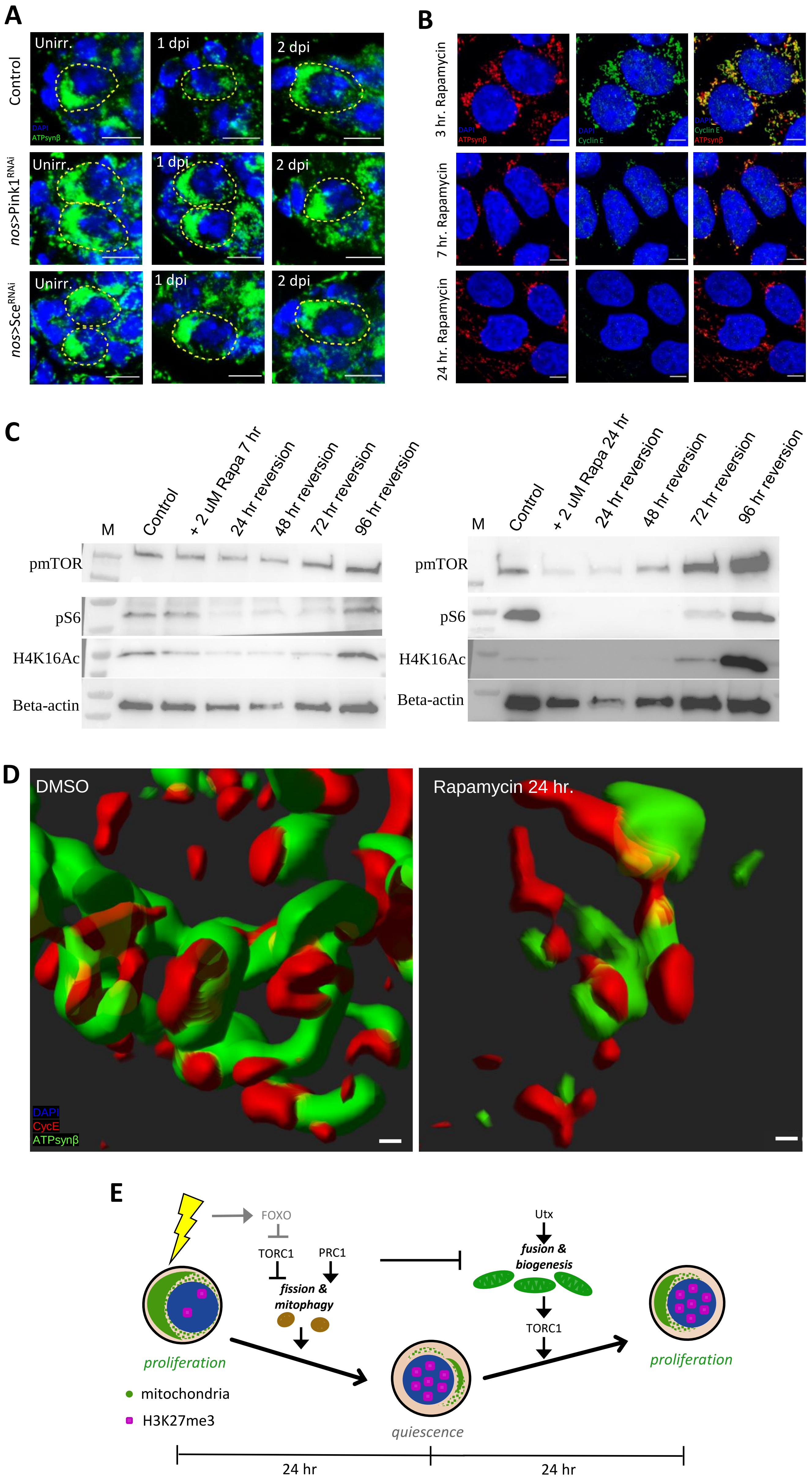

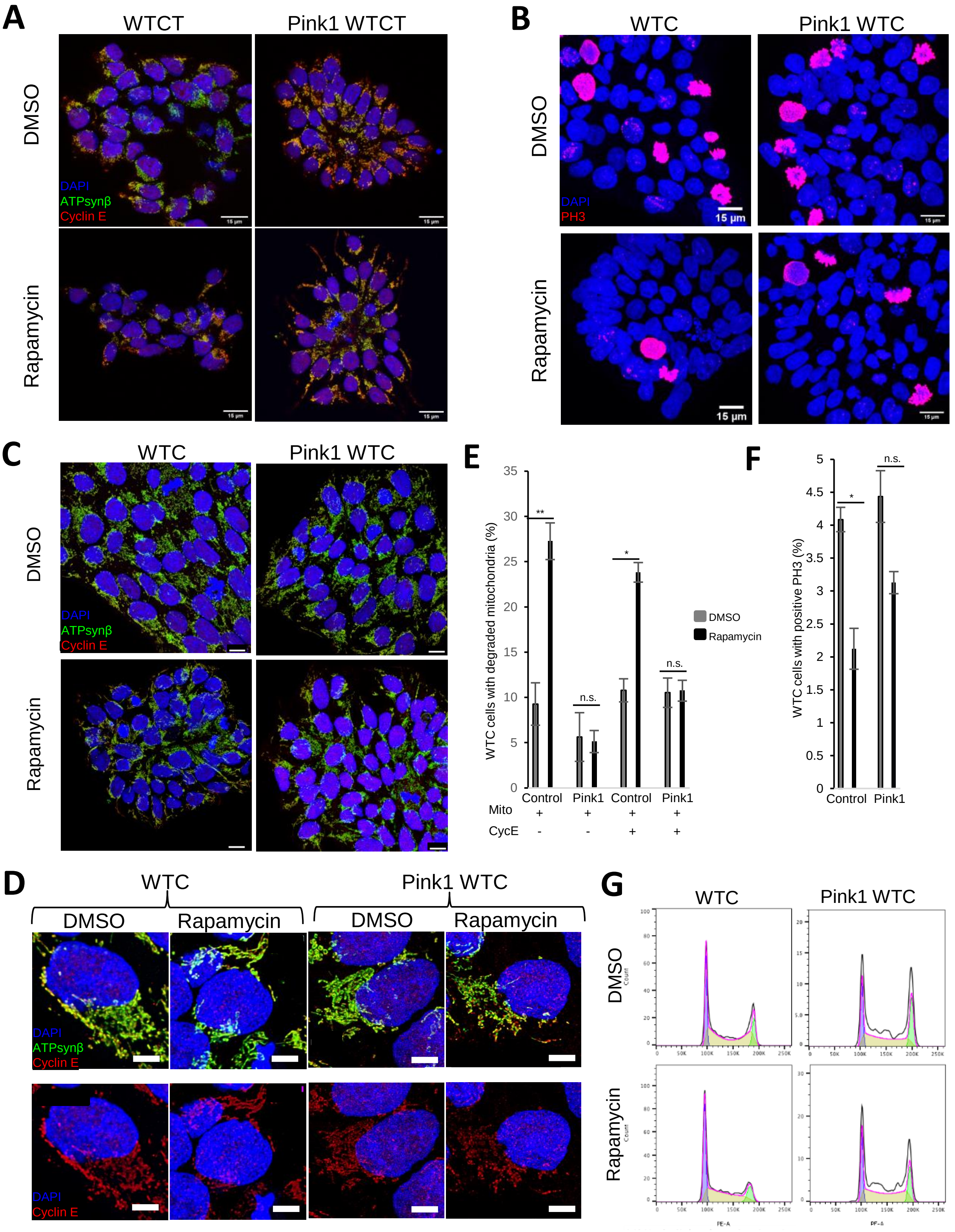
